# Allelic variations in a serine protease effector within *Clavibacter michiganensis* populations determine pathogen host range

**DOI:** 10.1101/2023.07.22.549466

**Authors:** Raj Kumar Verma, Veronica Roman-Reyna, Gitta L. Coaker, Jonathan M. Jacobs, Doron Teper

## Abstract

Plant pathogenic bacteria often have a narrow host range, which can vary among different isolates within a population. Here we investigated the host range of the tomato pathogen *Clavibacter michiganensis* (Cm). We determined the genome sequences of 40 tomato Cm isolates and screened them for pathogenicity on tomato and eggplant. Our screen revealed that out of the tested isolates, five were unable to cause disease on any of the hosts, 33 were exclusively pathogenic on tomato, and two were capable of infecting both tomato and eggplant. Through comparative genomic analyses, we identified that the five non-pathogenic isolates lacked the *chp/tomA* pathogenicity island, which has previously been associated with virulence in tomato. In addition, we found that the two eggplant-pathogenic isolates encode a unique allelic variant of the serine protease *chpG* (*chpG*^C^), an effector that is recognized in eggplant. Introduction of *chpG*^C^ into a *chpG* inactivation mutant in the eggplant-non-pathogenic strain Cm101, failed to complement the mutant, which retained its ability to cause disease in eggplant and failed to elicit hypersensitive response (HR). Conversely, introduction of the *chpG* variant from Cm101 into an eggplant pathogenic isolate (CmC48), eliminated its pathogenicity on eggplant, and enabled CmC48 to elicit HR. Our study demonstrates that allelic variation in the *chpG* effector gene is a key determinant of host range plasticity within Cm populations.

## Introduction

Many bacterial plant pathogens are specialists by nature and found in association with a small number of hosts (1). This phenomenon is reflected by the dichotomy of the broad host range of plant-associated bacteria such as *Xanthomonas* and *Pseudomonas syringae* on a genus or species level and the extremely narrow reported host range of pathovar or sub-species of the aforementioned groups (2)(3). The true host range of bacterial plant pathogens is usually not well defined, and, with the increased availability of sequencing technologies, both subspecies and pathovar classification and their reported host range are subjected to constant changes (4)(5). Compatibility with the unique physiology and genetic background of plant species or cultivar is one of the main features that determine whether a pathogen can utilize a plant as a host even when ecological factors are bypassed through artificial infection. Factors like the detoxification of plant-specific antimicrobials, the ability to degrade unique structural macromolecules, targeting host-specific susceptibility targets, and, avoidance or deactivation of host immune signaling have all been reported to play a key role in determining host specificity (6)(7)(8). The most extensively studied host-determining factor in plant-pathogen interactions is the gene-for-gene type of immune recognition that confer recognition of pathogen’ effectors by host resistance protein (9). Through this mechanism, a species or cultivar-specific immune receptor activates a strong immune response upon recognition of a unique secreted or translocated pathogen effector (10). However, because of its elegant simplicity, gene-for-gene-based resistance can be relatively easily overcome by pathogens through modifications or removal of recognized effectors (11)(12)(13).

Bacterial canker caused by the actinobacteria *Clavibacter michiganensis* (Cm) is one of the most destructive bacterial diseases of tomato (14). The disease is characterized by the appearance of stem cankers, wilting, ‘bird’s eye’ fruit lesions, vascular collapse, and, in severe cases, death (14). Cm enters the host through wounds and natural openings and extensively colonizes the host xylem vessels, spreading to all of the plant aerial tissues, including fruits and seeds (15)(16). Seed colonization is essential for the long-distance dispersal of Cm and its introduction into new habitats (17). Through introduction by contaminated seeds, Cm was spread into most tomato-growing regions around the world and is considered an endemic pest in many countries that significantly affect tomato production (14). Because of the economic significance of the disease and the high risk of reintroduction via contaminated seeds, Cm populations were subjected to numerous phylogenetic analyses. These studies identified that the diversity within the Cm populations varies between countries. Cm populations in Turkey, Italy, and central Chile demonstrated low diversity, suggesting they mainly originated from founder populations that were established endemically in these countries (18)(19)(20). On the other hand, high diversity was observed in Cm populations in Israel, New York State, Argentina, Iran, and Greece (21)(22)(23)(24)(25) suggesting multiple introductions of Cm, presumably through contaminated seeds.

Unlike many plant-pathogenic gram-negative bacteria, Cm does not possess a type III secretion system capable of delivering effectors directly inside hosts cells. Instead, Cm pathogenicity is heavily relies on secreted extracellular hydrolases and apoplastic effectors (26)(27)(28)(29). The main virulence determinants of Cm are encoded within three genomic regions: the pCM1 and pCM2 plasmids and the *chp/tomA* pathogenicity island (PAI), a 129 kb genomic island localized within the Cm chromosome (30). pCM2 and the *chp/tomA* PAI encode for numerous secreted serine peptidases that are classified into two main protein families, the Chp/Pat-1 proteases and the Ppa proteases. Both of these protein families play a crucial role in the virulence of Cm (31)(32)(33).

Similar to other plant–pathogenic *Clavibacter* sp., Cm is a specialized pathogen that harbors a narrow host range. In field conditions, Cm is almost solely found in tomato, which serves as its main host (34) while infection in other Solanaceous crops such as potato, pepper, and eggplant is seldom reported (19)(35). The true host range of Cm is unclear and artificial infections of alternative hosts such as pepper and eggplant produce unstable and even contradictory results when conducted by different groups (36)(37)(38). Recent studies by ourselves and Boyaci et al. 2021, identified that most domesticated eggplant varieties demonstrate moderate to high resistance to a number of Cm strains (39)(40). We identified that eggplant resistance to Cm is facilitated through immune recognition of ChpG, a secreted serine peptidase of the Chp/Pat-1 family encoded by the *chp/tomA* PAI (40). A *chpG* inactivation mutant in the background of the Cm model strain Cm^101^ cannot cause HR and is fully pathogenic in numerous eggplant varieties, indicating that ChpG is a recognized effector and, therefore, its recognition in eggplant is likely to follow the gene-for-gene model (40). However, it is unclear whether this phenomenon is conserved in other clones within Cm populations.

In this study, we used functional and genomic analyses to determine the virulence and host range of 40 representative Cm isolates and identified that allelic variations in the *chpG* effector acts as a host range determinant.

## Results

### *Clavibacter michiganensis* isolates demonstrate variation in pathogenicity and host range

Several studies from the past decade reported that a wide variety of *Clavibacter michiganensis* (Cm) isolates cannot cause disease in many eggplant accessions in controlled artificial inoculations (37)(39)(40). These reports contradict the European and Mediterranean Plant Protection Organization (EPPO) Cm information page (EPPO code CORBMI) and earlier studies from the 1970s (36). Considering that the majority of tested eggplant accessions demonstrated moderate to high resistance to Cm (39), we hypothesized that pathogenicity on eggplant is a unique feature of specific pathotypes within Cm populations. To test this, we screened a library of Cm isolates for virulence on eggplant and tomato. The Cm isolate library was composed of 40 isolates: 37 isolates were collected over a period of 29 years (from 1994 to 2023) from in various regions in Israel and three additional reference clones originated from USA (C30 and C31) and the Netherlands (C20) (Table S1). All isolates were tested for pathogenicity on tomato and eggplant by monitoring wilting (on tomato) or leaf blotch (on eggplant) symptoms and quantifying stem bacterial populations (Table 1, figures 1, S1 and S2). We used the Cm model strains Cm^101^ and Cm^101^Ω*chpG* as a reference. Cm^101^Ω*chpG* is a *chpG* inactivation mutant in the background of Cm^101^ which we previously reported to be pathogenic on eggplant varieties (40). 35 out of the 40 tested isolates were pathogenic on tomato, and were able to cause wilt symptoms and colonize tomato stems to approximately 10^8^ −10^10^ CFU/gram tissue (Table 1, figures 1 and S1).

**Figure 1.**
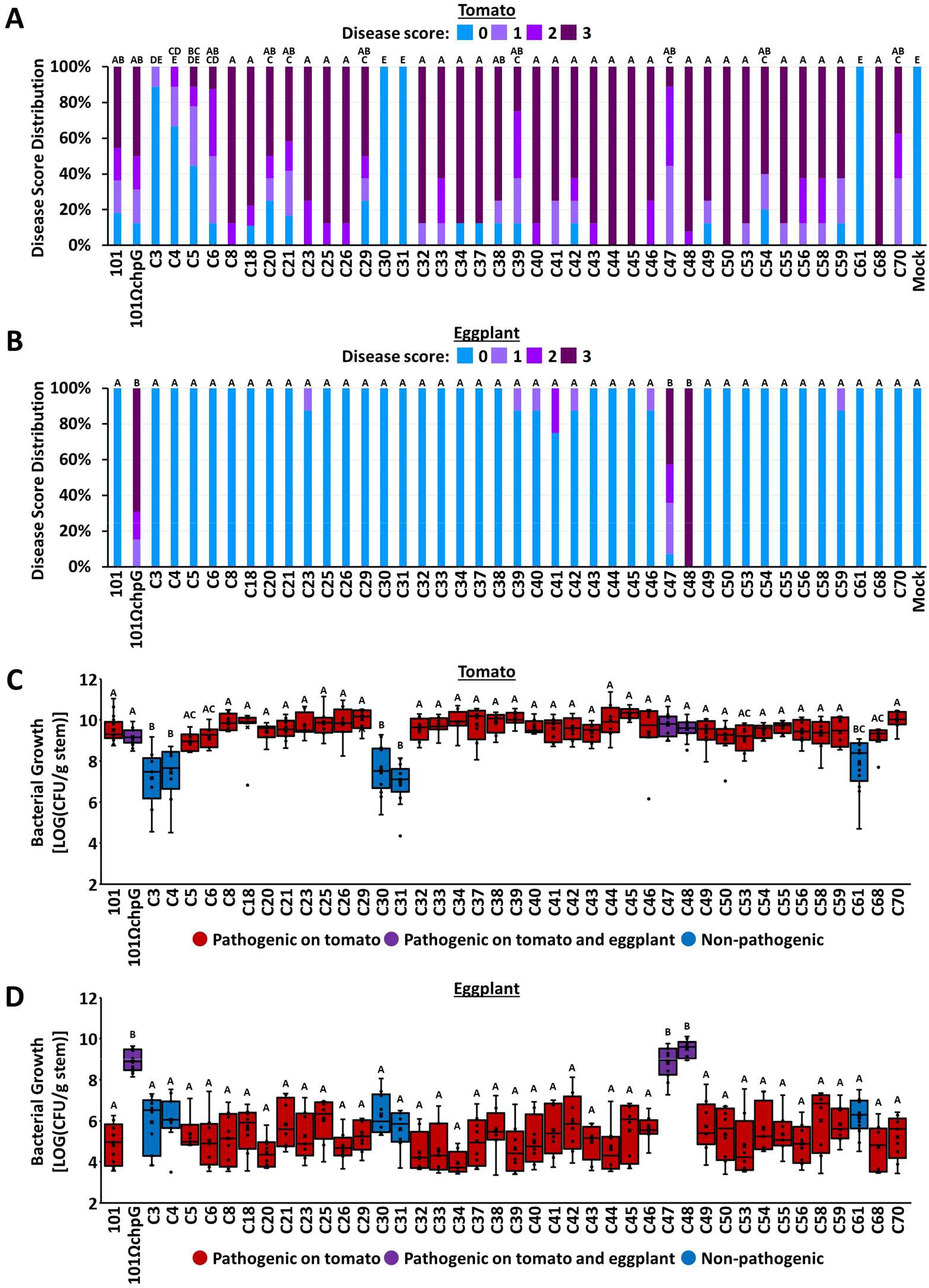
Symptoms and bacterial growth in plants inoculated with *Clavibacter michiganensis* (Cm) isolates. Four-leaf stage “Moneymaker” tomato plants (A, C) or three-leaf stage “Black Queen” eggplants (B, D) were inoculated with the indicated Cm isolates or water control (mock) by puncturing the stem area between the cotyledons with a wooden toothpick incubated in *Cm* solution (5 × 10^7^ CFU/ml). **(A**, **B**) Graphs represent the distribution of symptom severity in response to each isolate in at least eight plants taken from at least two experimental repeats. Symptoms severity were scored at 14 days post inoculations (dpi) according to the percentage of leaves displaying wilt (tomato) or leaf blotch (eggplant) symptoms by the following scale: 0 = no wilting/leaf blotch, 1 = 1-25%, 2 = 25-50%, 3 = 50-100%. **(C**, **D**) Stem bacterial populations 1 cm above the inoculation sites were quantified at 14 dpi. Lower and upper quartiles are marked at the margins of the boxes. Central lines, “×” and “o” represent medians, means and data points of at least eight biological repeats collected from at least two independent experiments. Boxes marked in red represent isolates that caused symptoms in tomato but not in eggplant, boxes marked in purple represent isolates that caused symptoms in tomato and in eggplant, and boxes marked in blue represent isolate that failed to cause symptoms in either tomato or eggplant. All depicted data were analyzed using one-way ANOVA followed by post-hoc Tukey HSD test. Letters indicate similarity in disease severity and bacterial populations (Tukey HSD test, *p value* ≤ 0.05).

**Table 1.** Virulence analysis of Cm isolates.

| Isolate | Tomato symptoms<sup>1</sup> | Tomato growth<sup>2</sup> | Eggplant symptoms<sup>1</sup> | Eggplant growth<sup>2</sup> | Eggplant HR | *chpG* variant<sup>3</sup> | *chp/tomA*<sup>4</sup> |
| --- | --- | --- | --- | --- | --- | --- | --- |
| Cm<sup>101</sup> | ++ | +++ | - | +/- | + | A<sup>S</sup> | + |
| Cm<sup>101</sup>Ω*chpG* | ++ | +++ | +++ | +++ | - | Ω<sup>P</sup> | + |
| C3 | +/- | + | - | +/- | - | -<sub>G,P</sub> | - |
| C4 | +/- | + | - | +/- | - | -<sub>G,P</sub> | - |
| C5 | + | +++ | - | +/- | + | B1<sup>G,S</sup> | + |
| C6 | ++ | +++ | - | +/- | + | D<sup>G,S</sup> | + |
| C8 | +++ | +++ | - | +/- | + | B2<sup>G</sup> | + |
| C18 | +++ | +++ | - | +/- | + | B1<sup>G,S</sup> | + |
| C20 | ++ | +++ | - | +/- | + | B1<sup>G</sup> | + |
| C21 | ++ | +++ | - | +/- | + | B1<sup>G,S</sup> | + |
| C23 | +++ | +++ | +/- | +/- | + | B1<sup>G</sup> | + |
| C25 | +++ | +++ | - | +/- | + | B1<sup>G</sup> | + |
| C26 | +++ | +++ | - | +/- | + | B1<sup>G</sup> | + |
| C29 | ++ | +++ | - | +/- | + | B2<sup>G,S</sup> | + |
| C30 | - | ++ | - | +/- | - | -<sub>G,P</sub> | - |
| C31 | - | + | - | +/- | - | -<sub>G,P</sub> | - |
| C32 | +++ | +++ | - | +/- | + | B1<sup>G</sup> | + |
| C33 | +++ | +++ | - | +/- | + | B1<sup>G</sup> | + |
| C34 | +++ | +++ | - | +/- | + | B1<sup>G</sup> | + |
| C37 | +++ | +++ | - | +/- | + | B1<sup>G</sup> | + |
| C38 | ++ | +++ | - | +/- | + | B1<sup>G</sup> | + |
| C39 | ++ | +++ | +/- | +/- | + | B1<sup>G</sup> | + |
| C40 | +++ | +++ | +/- | +/- | + | B1<sup>G</sup> | + |
| C41 | +++ | +++ | +/- | +/- | + | B1<sup>G</sup> | + |
| C42 | ++ | +++ | +/- | +/- | + | B1<sup>G</sup> | + |
| C43 | +++ | +++ | - | +/- | + | B1<sup>G</sup> | + |
| C44 | +++ | +++ | - | +/- | + | B1<sup>G</sup> | + |
| C45 | +++ | +++ | - | +/- | + | B1<sup>G,S</sup> | + |
| C46 | +++ | +++ | +/- | +/- | + | B1<sup>G,S</sup> | + |
| C47 | ++ | +++ | ++ | +++ | - | C<sup>G,S</sup> | + |
| C48 | +++ | +++ | +++ | +++ | - | C<sup>G,S</sup> | + |
| C49 | ++ | +++ | - | +/- | + | B1<sup>G</sup> | + |
| C50 | +++ | +++ | - | +/- | + | B1<sup>G</sup> | + |
| C53 | +++ | +++ | - | +/- | + | B1<sup>G</sup> | + |
| C54 | ++ | +++ | - | +/- | + | B1<sup>G</sup> | + |
| C55 | +++ | +++ | - | +/- | + | B1<sup>G</sup> | + |
| C56 | +++ | +++ | - | +/- | + | B1<sup>G</sup> | + |
| C58 | +++ | +++ | - | +/- | + | B2<sup>G</sup> | + |
| C59 | ++ | +++ | +/- | +/- | + | B2<sup>G</sup> | + |
| C61 | - | ++ | - | +/- | - | -<sup>G,P</sup> | - |
| C68 | +++ | +++ | - | +/- | + | B1<sup>G,S</sup> | + |
| C70 | ++ | +++ | - | +/- | + | B1<sup>G</sup> | + |

<sup>1</sup>Average symptoms at 14 days post inoculation. (-) = no symptoms. (+/-) = symptom score between 0.1-1, (+) = symptom score between 1-1.5, (++) symptom score between 1.5-2.5, (+++) = symptom score between 2.5-3.
<sup>2</sup>Average bacterial growth (BG) [LOG (CFU/gram stem)] at 14 days post inoculation. (+/-) = $BG \leq 6$ , (+) = $6 < BG \leq 7.5$ , $BG \leq 6$ , (++) = $7.5 < BG \leq 8.5$ , (+++) = $8.5 < BG$ .
<sup>3</sup>Allelic variant of *chpG* according to genome sequencing (G) and/or PCR (P) following Sanger sequencing (S).
<sup>4</sup>Presence of the *chp/tomA* genomic island according to genome sequencing. NT – genome is not sequenced.

However, high variations were observed in the intensity of wilt symptoms and colonization capacity between the different pathogenic isolates (Figure 1). Five isolates (i.e. C3, C4, C30, C31, and C61) were non-pathogenic on tomato. These isolates failed to cause wilt symptoms and demonstrated a significant reduction in tomato stem colonization compared to the other clones (Table 1, figures 1 and S1). In contrast to the tomato assays, 38 out of the 40 tested Cm isolates failed to cause leaf blotch symptoms on eggplant and were only able to colonize eggplant stems to approximately 10^4^ −10^6^ CFU/gram tissue (Table 1, figures 1 and S2). Two isolates, C47 and C48, were fully pathogenic on eggplant. These isolates caused leaf blotch symptoms and colonize eggplant stems 100-1000 fold higher than the other tested isolates, reaching approximately 10^8^ −10^9^ CFU/gram tissue (Table 1, figures 1 and S2).

We previously reported that eggplant resistance to the Cm model strain Cm^101^ is accompanied by activation of hypersensitive response (HR) (40). Hence, we monitored whether the Cm isolates in our library triggered HR in eggplant, and discovered that 33 isolates elicited HR in eggplant leaves. These include all tomato-pathogenic isolates with the exception of the two eggplant-pathogenic isolates C47 and C48 (Table 1, figure 2). Surprisingly, all five tomato-non-pathogenic isolates (C3, C4, C30, C31, and C61) failed to elicit HR in eggplant as well, suggesting they lack a HR-inducing elicitor that is found in the tomato-pathogenic clones (Table 1, figure 2). Our screen demonstrated that Cm pathogenicity on eggplant is isolate-dependent and associated with the inability to elicit HR on eggplant leaves.

**Figure 2.**
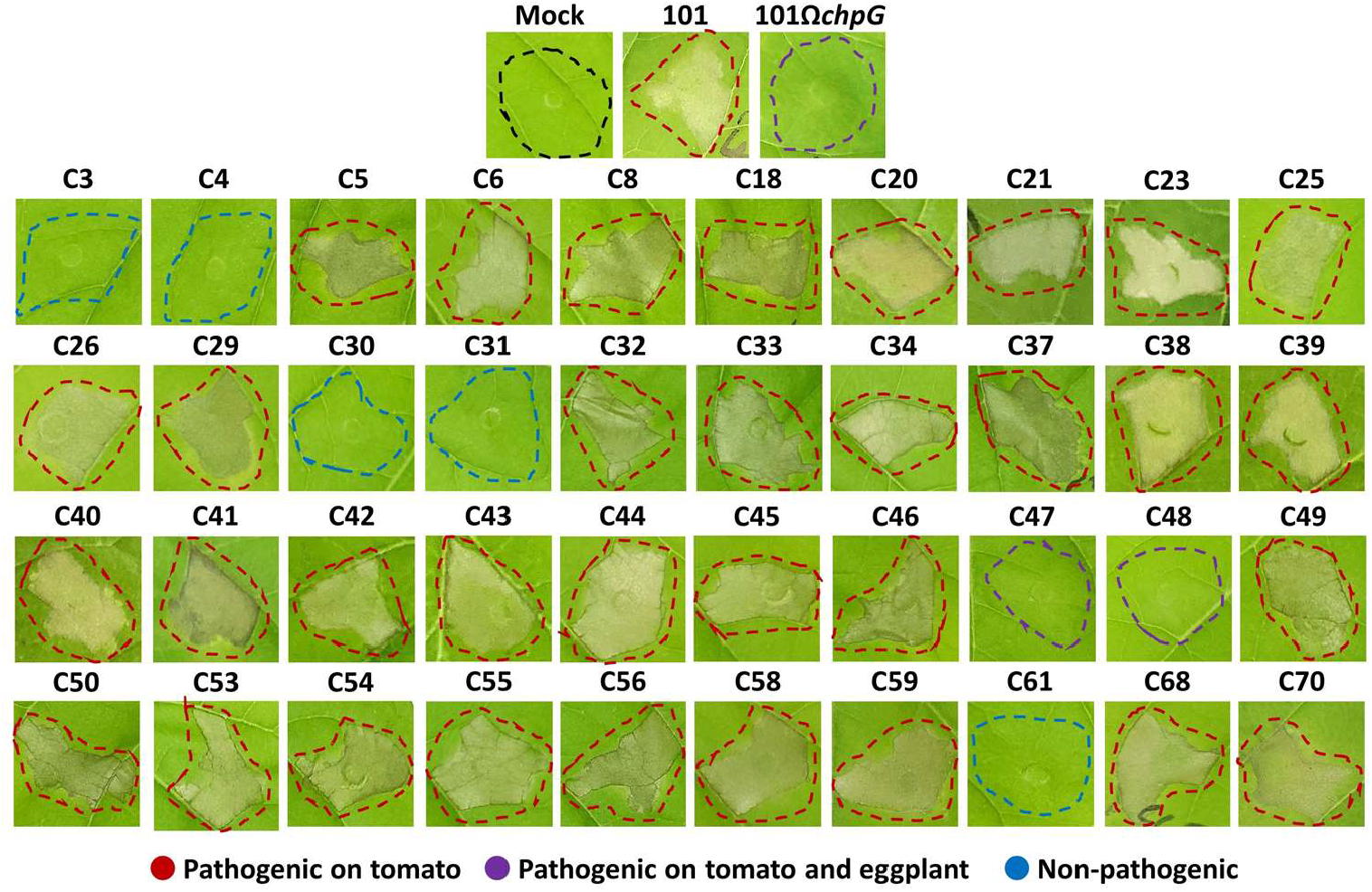
Elicitation of the hypersensitive response (HR) in eggplant leaves by *Clavibacter michiganensis* (Cm) isolates. Five to six leaf stage “Black Queen” eggplant leaves were infiltrated (10^8^ CFU/ml) with the indicated Cm isolates. Pictures were taken 48 h post infiltration. Infiltrated area are marked in colored dotted lines. Red lines represent isolates that caused symptoms in tomato but not in eggplant, purple lines represent isolates that caused symptoms in tomato and in eggplant, and blue lines represent isolate that failed to cause symptoms in either tomato or eggplant. Pictures are representatives of one out of at least 10 repeats from at least two independent experiments.

### Genomic analysis of Israeli Cm isolates

The Cm isolates used in our screen demonstrated high variation in their virulence and host range. To identify the source of these variations, we determined the genome sequences of the isolates in our library using the Illumina Nextseq2000 platform (BioProject ID PRJNA966807)(Table S2). Next, we determined the phylogenetic lineage of the Cm isolates by conducting multiple sequence alignment of core genes (Figure 3A) using the M1CR0B1AL1Z3R web server (41). The analysis was conducted along with NCBI genome deposits of Cm and non-Cm *Clavibacter* genomes, which were used as reference points. As expected, all 40 isolates clustered together with the NCBI Cm genome deposits (Figure 3A). These also include the eggplant-pathogenic and tomato non-pathogenic clones, indicating that virulence and host range variations exist within Cm populations. In addition, we observed multiple distinct phyletic clusters within the Israeli Cm isolates, some of which also included non-Israeli reference genomes. This observation supports previous fingerprinting-based phylogenetic analysis of Cm populations in Israel conducted in our institute (21)(42)(Table S1) and suggests that Cm populations originated from multiple independent introductions that occurred throughout the years and not from a local founder population. While both eggplant-pathogenic isolates, C47 and C48, cluster together in the same lineage (Figure 3A, marked in purple), the five tomato-non-pathogenic isolates cluster into three independent lineages (Figure 3A, marked in blue), two of which contained tomato-pathogenic clones as well.

**Figure 3.**
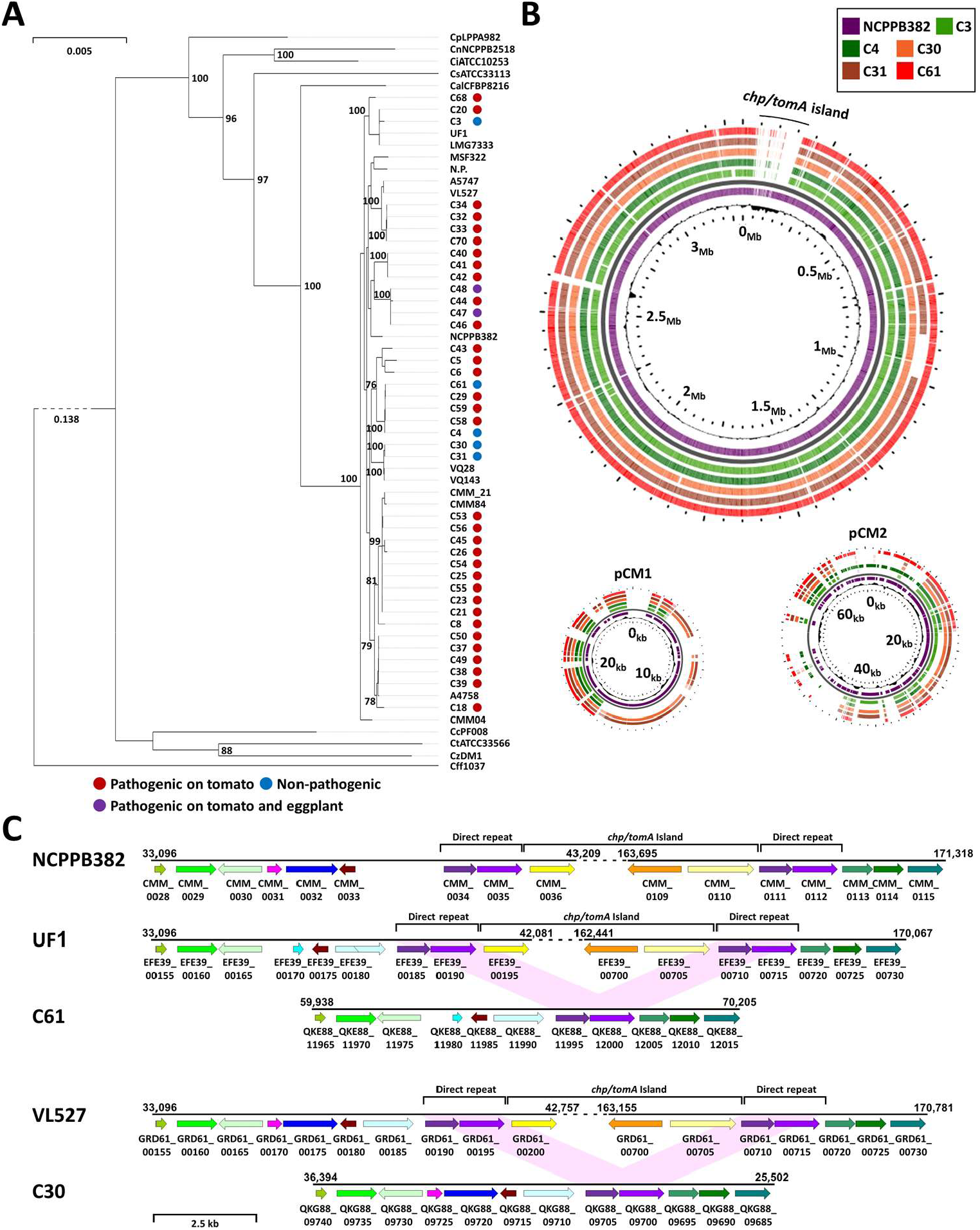
Tomato non-pathogenic *Clavibacter michiganensis* (Cm) isolates lack the *chp/tomA* pathogenicity island. (**A**) Phylogenetic tree of Cm isolates used in this study. The tree was produced using the M1CR0B1AL1Z3R web server (https://microbializer.tau.ac.il/) and is based on maximum-likelihood multiple sequence alignments of 129 core genes under default features and visualized by PhyD3. Genomes used for the analysis are 40 genomes of Cm isolates characterized in this study; references Cm strains UF1, LMG7333, NCPPB382, CMM84, Cmm_21, VQ143, VQ28, VL527, A5747, N.P., MSF322, and CMM04; and references strains of *C. sepedonicus* ATCC331133, *C. capsici* PF008, *C. nebraskensis* NCPPB2581*, C. insidiosus* ATCC10253*, C. tessellarius* ATCC33566*, C. zhangzhiyongii* DM1*, C. phaseoli* LPPA982, and *C. californiensis* CFBP8216. *Curtobacterium flaccumfaciens* Cff1037 was used as an outgroup. Isolates which were pathogenic on tomato but not pathogenic on eggplant are marked with a red dot, isolates which were pathogenic on tomato and eggplant are marked with a purple dot, and isolates which were non-pathogenic on tomato and eggplant are marked with a blue dot. (**B**) Whole genome alignment of the tomato non-pathogenic Cm isolates C3, C4, C30, C31, and C61 was performed against CDS of Cm strain NCPPB382 chromosome (NCBI GenBank: AM711867), pCM1 plasmid (AM711865) and pCM2 plasmid (AM711866) and visualized with BLAST atlas analysis in Gview server (https://server.gview.ca/) using default features. The *chp/tomA* genomic island (positions 42,081-162,441 in the NCPPB382 chromosome) is labeled. (**C**) Physical maps of the area surrounding the *chp/tomA* island in the tomato-pathogenic reference strains NCPPB382 (AM711867), UF1 (NZ_CP033724), and VL527 (NZ_CP047054) and representative corresponding regions in the tomato non-pathogenic isolates C61 [SAMN34566112, contig 16. Similar gene synteny was observed in C3 (SAMN34566069, contig 13, 59,987 – 70,253) and C4 (SAMN34566070, contig 14, 59,985 – 70,252)], and C30 [SAMN34566083, contig 9. Similar gene synteny was observed in C31 (SAMN34566084, contig 9, 36,376 – 25,484)]. Predicted ORFs are marked in arrows supplemented with locus tags in the corresponding genomes. Similar colors indicates the ORFs share DNA sequence identify of >97%, “/” indicates an ORF was disrupted by a frameshift mutation. The *chp/tomA* and direct repeat region are marked. The first and two last ORFs of the *chp/tomA* region are labeled in shades of yellow. The two ORFs encoded within the direct repeat region are labeled in shades of purple.

To assess the genetic differences between our isolates we conducted comparative genomic analysis by utilizing BLAST atlas feature in the Gview server (43). All analyses were done in comparison to the Cm model strain NCPPB382 that served as a reference point. The comparative analysis identified that the *chp/tomA* pathogenicity island (PAI), which was reported to be the major chromosomal-encoded genomic region associated with virulence (30) was absent in all five tomato-non-pathogenic strains (Figure 3B). In contrast, the *chp/tomA* PAI was present in all sequenced tomato-pathogenic isolates (Figure S3). In addition, the presence of ORFs associated with the pCM1 and pCM2 plasmids demonstrated high variability between isolates (Figure S3). Most plasmid variations were in genes that were reported to be required for plasmid maintenance (44)(45) while genes associated with virulence such as *celA* and *pat-1* were conserved in most isolates (Figure S3). This suggests that some isolates harbor plasmids with different replicons than that of pCM1 and pCM2 or potential genomic integration of the virulence-associated functions of the plasmids into the chromosome.

The absence of the *chp/tomA* PAI in the tomato non-pathogenic isolates can be an indicator that these isolates are either ancestral *Clavibacter* species that have yet to acquire the *chp/tomA* PAI or that the *chp/tomA* PAI was lost in the course of evolution. The scattered phyletic distribution of the tomato-non-pathogenic isolates and their phylogenetic clustering along with tomato-pathogenic isolates suggests a loss of the PAI. In further inquiry, we took a closer look at the surroundings of the region that the *chp/tomA* PAI was supposedly initially integrated into in both pathogenic and non-pathogenic clones. As previously reported, in many pathogenic isolates, the *chp/tomA* regions were flanked by two ∼1.9 kb direct repeats that share 98-99% DNA sequence identity (30)(46). These direct repeats match to positions 40,054-41,981 and 166,885-168,812 in the chromosome of Cm model strain NCPPB382 (Figures 3C and S4, marked in purple). Interestingly, all five tomato-non-pathogenic isolates harbored a single copy of this region that was surrounded in homologous areas to the upstream and downstream regions to the *chp/tomA* PAI in the pathogenic isolates (Figures 3C and S4). This suggest that the *chp/tomA* PAI was lost due to potential recombination between the direct repeats and that this event is more likely to occur independently in different isolates.

### Eggplant-pathogenic Cm isolates encode a unique allelic variant of the *chpG* effector

We previously reported that eggplant resistance to the Cm model strain Cm^101^ is mediated through HR-based immune recognition of the secreted serine protease ChpG (40). Therefore, we hypothesized that eggplant-pathogenic and HR-negative isolates are able to evade host recognition because they either lack a functional *chpG* homolog or encode for a *chpG* variant that is not recognized in eggplants. To examine this, we conducted *in silico* analysis for occurrence and polymorphisms in the *chpG* gene in our Cm isolate library. Our analysis identified five *chpG* allelic variants within Cm populations (Table 1) which we named *chpG*^A^, *chpG*^B1^, *chpG*^B2^,*chpG*^C^ and *chpG^D^*(Figures S5 and S6).

The *chpG*^A^ variant (CMM_0059) is found solely in the model strains NCPPB382 and Cm^101^. The *chpG*^B1^ variant (EFE39_00385) is the dominant variant in our library and the available NCBI deposits (table 1).The *chpG*^B2^ variant (LHJ47_00410) is found in four of the isolates from our library (table 1) and the NCBI Cm deposits CMM39 and CMM04. *chpG*^C^ and *chpG*^D^ variants were unique to isolates in our library and were not identified in any of the NCBI Cm genome deposits. The *chpG*^C^ variant (QKF70_15430) was only present in the two eggplant-pathogenic isolates C47 and C48 while the *chpG*^D^ variant (QKG63_14905) was present in isolate C6 (Table 1).

We next examined how these allelic variations translate to differences in amino acid (Figures 4A and S6) as compared to the most abundant *chpG* allelic variant *chpG*^B1^. *chpG*^B2^ harbor a C78->T nucleotide substitution that results in a synonymous mutation and therefore did not affect amino acid composition, which was identical to that of *chpG*^B1^. We defined this protein variant as ChpG^B^. *chpG*^A^ has a single T65->C substitution that results in L22->P amino acid alteration. *chpG*^C^ has a single T506->G substitution which results in V169->G amino acid alteration. Finally, *chpG*^D^ harbored multiple nucleotide substitutions and insertion, which resulted in several amino acid alterations including the insertion of N at amino acid position 142 and the amino acid substitutions of P147->L148 and G157->S158. Next, we mapped the amino acid modifications to the AlphaFold ChpG protein structure model (UniPort num’ A5CLZ4, Figure 4B)(47). ChpG is predicted to contain an unstructured N-terminal domain (positions 1-43) containing the sec-dependent signal peptide and a C-terminal serine protease domain composed of two beta-barrel domains which match the structure of other serine proteases (48) and an external beta-sheet region (positions 126-155) that expend outside of the main serine protease domain (Figures 4A and 4B). A similar structure was observed in other Chp proteases (*i.e.* Pat-1, ChpC, ChpF, ChpE, data not shown) indicating its significance to the function of this protein family. We mapped the predicted amino acid polymorphic sites into the ChpG structure (Figure 4B). The ChpG^A^ polymorphic site at position 22 is placed within the unstructured N-terminal signal sequence, the ChpG^C^ polymorphic site at position 169 is located within the main conserved serine protease domain beta-barrel structure while the three ChpG^D^ polymorphic sites are located within or in the proximity of the predicted external beta-sheet region (Figure 4B).

**Figure 4.**
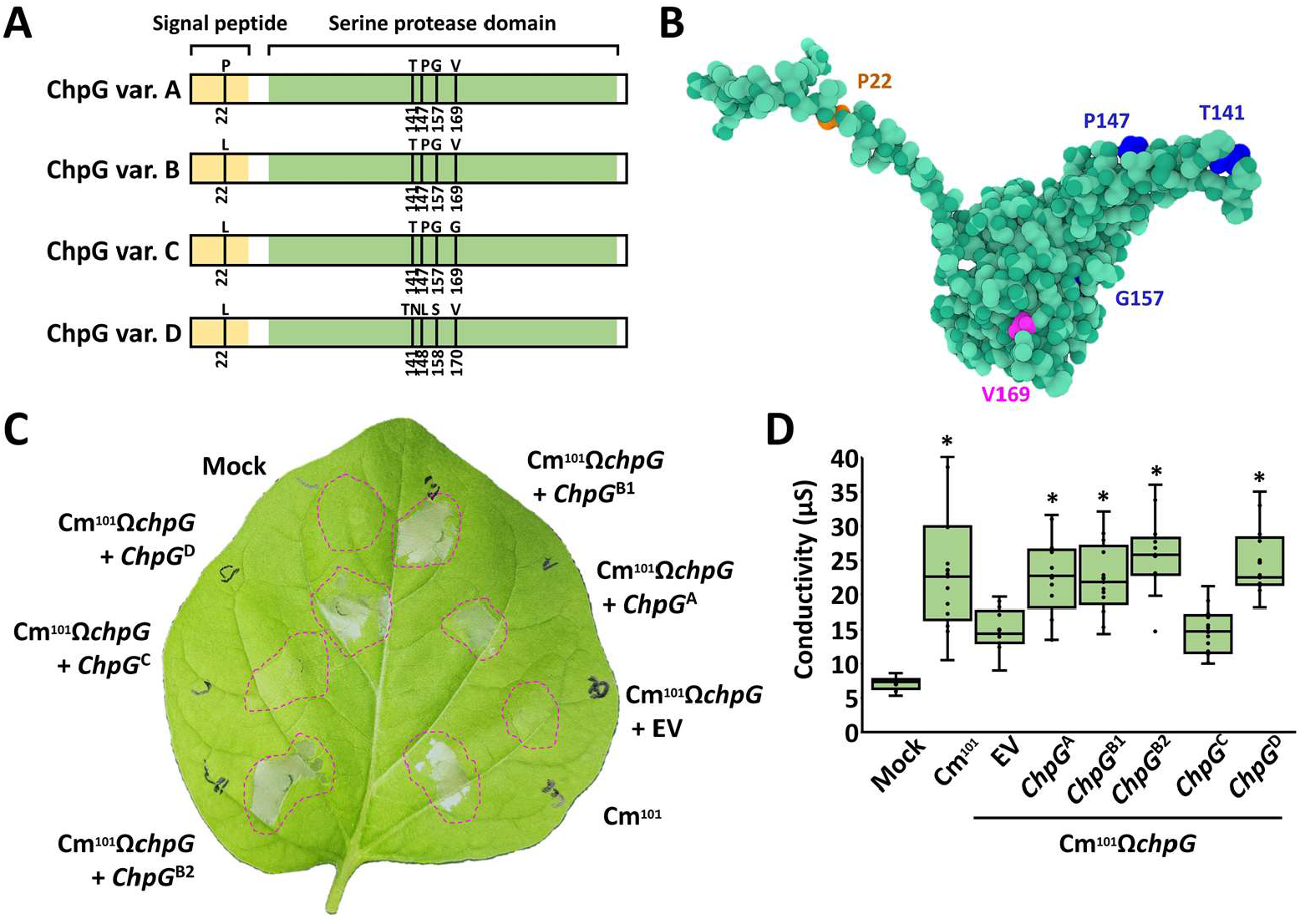
The ChpG allelic variants are differentially recognized in eggplant. **(A)** Schematic representation of the four ChpG protein variants depicted in figure S6. Signal peptide region, predicted by SignalP-5.0 (https://services.healthtech.dtu.dk/services/SignalP-5.0/) is labeled in yellow. Alpha-lytic serine protease domain, predicted by NCBI conserved domain search (https://www.ncbi.nlm.nih.gov/Structure/cdd/wrpsb.cgi) is labeled in green. Amino acid polymorphic sites are marked in black lines. **(B**) Structure prediction of ChpG variant A (ChpG^A^) was conducted by Alphafold (https://alphafold.ebi.ac.uk/entry/A5CLZ4, UniPort num’ A5CLZ4) and visualized by Mol* (https://molstar.org/viewer/). Amino acid polymorphic sites unique to ChpG^A^, ChpG^C^ and ChpG^D^ are respectively marked in orange, magenta and blue. (**C, D**) Black Queen eggplant leaves were infiltrated with 10 mM MgCl_2_ (mock) or suspensions (10^8^ CFU/ml) of Cm^101^, and Cm^101^Ω*chpG* clones expressing the indicated *chpG* variants or empty vector control (EV). (**C**) Representative picture was taken 48 h post infiltration (hpi). (**D**) Cell death was quantified by ion leakage at 36 hpi. Lower and upper quartiles are marked at the margins of the boxes. Central lines and “o” represent medians and data points of 21 biological repeats collected from three independent experiments. “*” indicates significant differences (Mann–Whitney U test, *p-value* ≤ 0.05) from Cm^101^Ω*chpG* + EV.

The *chpG*^C^ variant is uniquely encoded in the two eggplant-pathogenic strains C47 and C48, which suggests that this allelic variant is not recognized in eggplant. To test this, we monitored whether each one of the five *chpG* allelic variants can complement Cm^101^Ω*chpG* and restore recognition. All five *chpG* allelic variants were fused to a triple HA tag, cloned into the *E. coli*-*Clavibacter* shuttle vector pHN216 under the control of the constitutive *pCM1* promoter and introduced into Cm^101^Ω*chpG*. Protein accumulation was confirmed in all transformants using western blot (Figure S7A). Eggplant leaves were infiltrated with cultures of Cm^101^Ω*chpG* carrying each of the allelic variants or empty vector control (EV) and monitored for HR for 48 h. HR was observed in Cm^101^Ω*chpG* carrying *chpG* variants A, B1, B2 and D while no HR was observed Cm^101^Ω*chpG* carrying *chpG* variant C (Figures 4C and 4D). Our data suggest that eggplant-pathogenic isolates harbor an adaptive modification in the *chpG* effector gene that abolish it’s recognition in eggplant and enable these isolates to evade activation of host immune response.

### Purified ChpG^C^ variant does not elicit HR in eggplant

We previously showed that purified ChpG protein cloned from the eggplant-non-pathogenic Cm strain Cm^101^ elicit host-specific HR in eggplants (40). We tested whether the ability to elicit HR is abolished in the purified ChpG^C^ variant, which failed to complement Cm^101^Ω*chpG*. The mature proteins of ChpG^AB^ (represent both ChpG^A^ and ChpG^B^ since mature variants lack the predicted secretion signal) and ChpG^C^ were fused to maltose-binding-protein tag, purified from *E. coli* (Figure 5A) and assayed for their ability to elicit HR in eggplant leaves upon syringe infiltration. The purified ChpG^AB^ variant, which is found in eggplant-non-pathogenic isolates, elicited strong HR in eggplant leaves between 36-48 h post infiltration (hpi) (Figures 5B and 5C) while the purified ChpG^C^ variant failed to do the same (Figures 5B and 5C). Our analysis confirms that ChpG^C^ variant is not recognized in eggplant.

**Figure 5.**
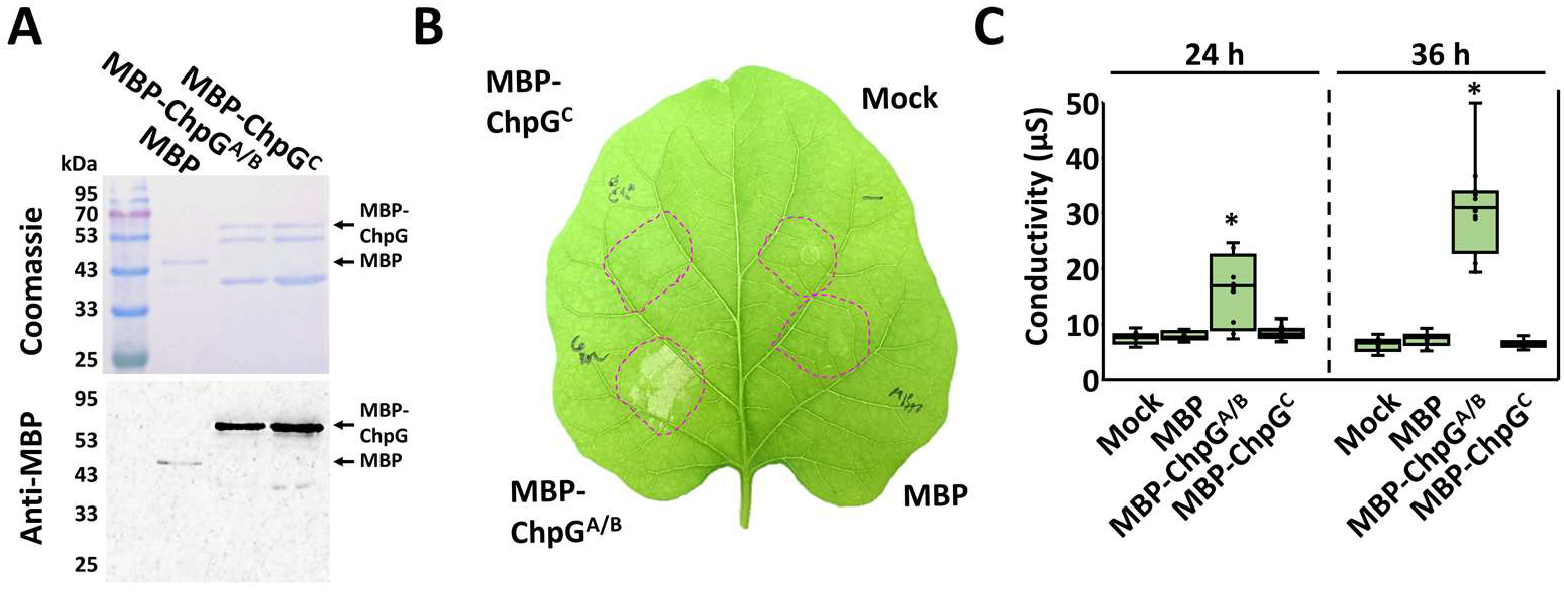
The ChpG^C^ variant does not elicit HR in eggplant. (**A**) Mature ChpG variants, lacking the predicted secretion sequence, were fused to a maltose-binding protein (MBP) tag, purified from *E. coli*, and visualized by SDS-PAGE using Coomassie blue staining (upper panel) and western blot analysis using anti MBP antibody (lower panel). **(B, C**) Purified proteins (0.01 µg/ml) or MgCl_2_ control (mock) were infiltrated into “Black Queen” eggplant leaves. **(B**) Representative leaf was photographed 36 hours post infiltration (hpi). (**C**) Cell death was quantified by ion leakage at 24 and 36 hpi. Lower and upper quartiles are marked at the margins of the boxes. Central lines and “o” represent medians and data points of at least 12 biological repeats collected from three independent experiments. “*” indicates a significant difference (Mann–Whitney U test, *p-value* ≤ 0.05) from mock control.

### Complementation analyses demonstrate differential recognition of *chpG* allelic variants

To prove that the occurrence of different allelic variants of *chpG* indeed determines the host range of Cm isolates, we conducted reciprocated complementation analyses. For that, pHN216 carrying *chpG*^A^ and *chpG*^C^ were introduced into the eggplant pathogenic isolate Cm C48 and Cm^101^Ω*chpG*, and transformants were assayed from HR elicitation and virulence in eggplant. Protein accumulation of ChpG^A^ and ChpG^C^ was monitored in the two strains and demonstrated that while both ChpG variants were present in a premature and mature forms, ChpG^A^ accumulation was significantly higher in both C48 and Cm^101^Ω*chpG* compared to ChpG^C^, suggesting that allelic polymorphism in *chpG* might affect its translation, secretion efficiency, or protein stability (Figures S7A, S7B and S7C). C48 and Cm^101^Ω*chpG* carrying pHN216:*chpG*^A^ were able to elicit HR upon infiltration into eggplant leaves between 24-48 hpi while C48 and Cm^101^Ω*chpG* carrying pHN216:*chpG*^C^ or pHN216 empty vector control failed to do the same (Figure 6A and 6B). Virulence assays were conducted using stab inoculation of eggplant stems. Virulence was quantified by monitoring leaf blotch symptoms and stem bacterial populations two weeks post inoculations. As previously reported, C48 and Cm^101^Ω*chpG* caused significant leaf blotch symptoms on eggplant (Figures 6C, 6D and 6E). However, introduction of pHN216:*chpG*^A^ but not pHN216:*chpG*^C^ turned both Cm isolates to non-pathogenic on eggplant (Figures 6C, 6D and 6E). Stem bacterial populations of C48 and Cm^101^Ω*chpG* carrying pHN216:*chpG*^C^ was similar to C48 and Cm^101^Ω*chpG* and reached approximately 10^9^ CFU/gram stem (Figure 5F). In contrast, bacterial population of C48 and Cm^101^Ω*chpG* carrying pHN216:*chpG*^A^ demonstrated 200-1000 fold reduction compared to their parental clones and reached approximately 10^6^ - 5 × 10^6^ CFU/gram stem (Figure 6F). To eliminate the possibility that introduction of pHN216:*chpG*^A^ or pHN216:*chpG*^C^ resulted in a non-host-specific alteration in virulence, C48 and Cm^101^Ω*chpG* transformed clones were assayed for virulence on tomato and demonstrated no significant changes in their ability to cause disease (Figure S8). Our data shows that the *chpG*^A^ significantly hinders the virulence of C48 and Cm^101^Ω*chpG* on eggplant while *chpG*^C^ does not.

**Figure 6.**
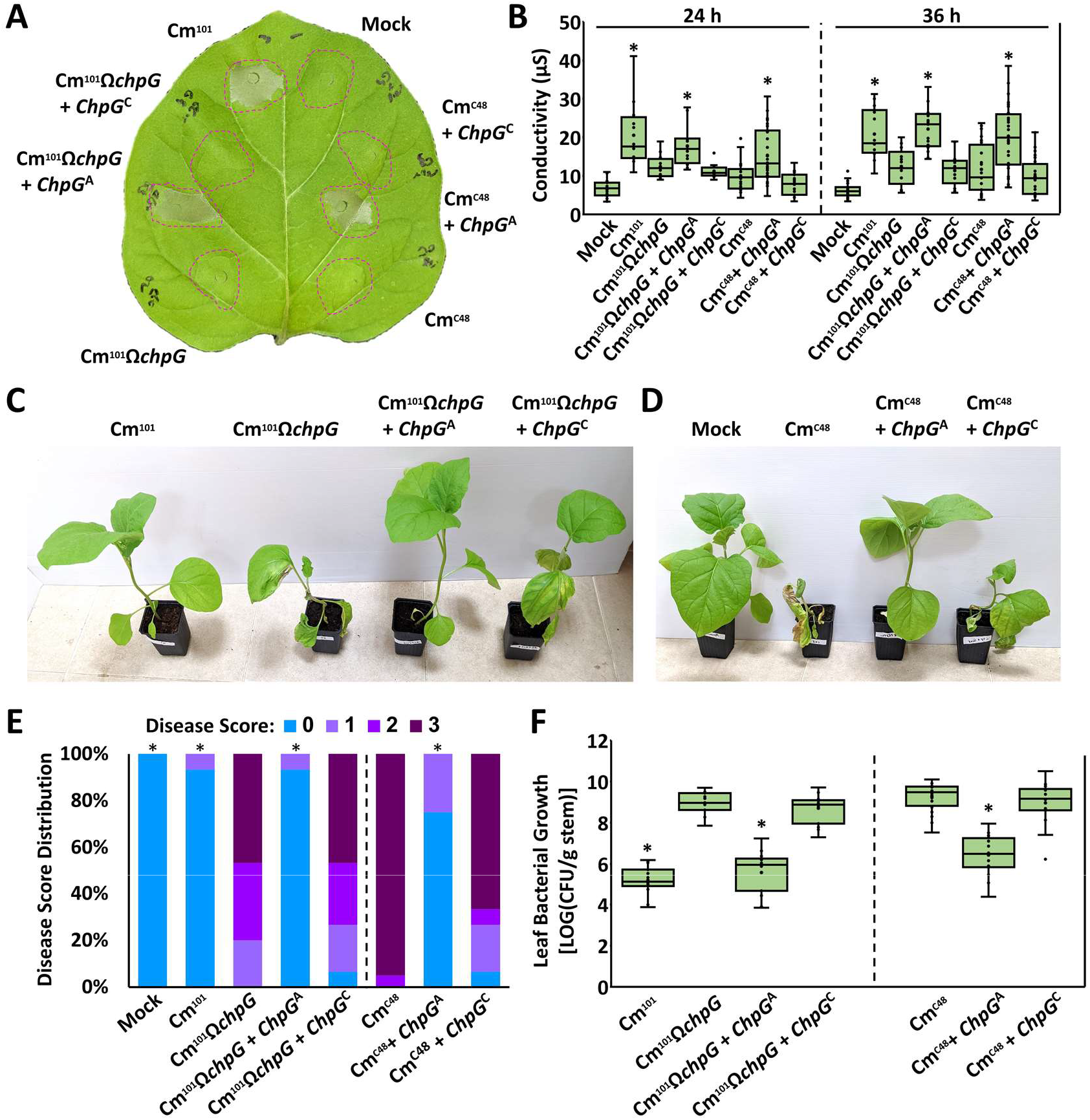
The ChpG^C^ variant does not restrict *Clavibacter michiganensis* (Cm) from colonizing eggplant. (**A, B**) Five to six leaf stage “Black Queen” eggplant leaves were infiltrated (10^8^ CFU/ml) with Cm^101^, Cm^101^Ω*chpG* and Cm^C48^ expressing the indicated *ChpG* variants or empty vector control (EV). (**A**) Picture was taken at 36 hours post inoculations (hpi). (**B**) Cell death was quantified by ion leakage at 24 and 36 hpi. Lower and upper quartiles are marked at the margins of the boxes. Central lines and “o” represent medians and data points of at least 12 (24 h) or 17 (36 h) biological repeats collected from at least two (24 h) or three (36 h) independent experiments. “*” indicates a significant difference (Mann– Whitney U test, *p-value* ≤ 0.05) from Cm^101^Ω*chpG* (left panel) or Cm^C48^ (right panel). (**C, D, E, F**) Three-leaf stage “Black Queen” eggplants were inoculated with the indicated Cm strains or water control (mock) by puncturing the stem area between the cotyledons with a wooden toothpick incubated in Cm solution (5 × 10^7^ CFU/ml). (**C, D**) Representative plants were photographed 14 days post inoculation (dpi). (**E**) Leaf blotch symptoms were quantified at 14 dpi according to the following scale: 0 = no leaf blotch, 1 = 1-25%, 2 = 25-50%, 3 = 50-100%. Graph depicts the symptom distribution in at least 15 plants pooled from at least three independent experiments. “*” indicates the score distribution is different from Cm^101^Ω*chpG* (left panel) or Cm^C48^ (right panel) (Pearson’s chi-squared test, *p-value* ≤ 0.05). (**F**) Stem bacterial populations 1 cm above the inoculation sites were quantified at 14 dpi. Lower and upper quartiles are marked at the margins of the boxes. Central lines and “o” represent medians and data points of 15 biological repeats collected from at least three independent experiments. “*” indicates a significant difference (Mann–Whitney U test, *p-value* ≤ 0.05) from Cm^101^Ω*chpG* (left panel) or Cm^C48^ (right panel).

### A single amino acid substitution in the ChpG serine protease domain eliminates its recognition in eggplant

ChpG^A^ and ChpG^C^ differ from each other due to two amino acid substitutions, at position 22 which is predicted to be part of the signal peptide and at position 169 that is part of the serine protease domain (Figure 4A). Purified mature ChpG^AB^, which lacks the signal peptide region, is sufficient to elicit HR in eggplant leaves (Figures 5B and 5C), which suggest that valine to glycine alteration at position 169 in ChpG^C^ allows it to evade recognition in eggplant. To test this, we conducted reciprocal substitutions of amino acid position 169 in ChpG^A^ and ChpG^C^. Val169 in ChpG^A^ was substituted to Gly (ChpG^A^_V169G_) while Gly169 in ChpG^C^ was substituted to Val (ChpG^C^_G169V_). We note that Leucine is found at position 22 in both ChpG^B^ and ChpG^C^ and therefore the amino acid sequence of ChpG^C^_G169V_ is identical to ChpG^B^. pHN216-based plasmids carrying ChpG^A^ or ChpG^C^_G169V_ fused to HA tag were introduced into Cm^101^Ω*chpG*. We monitored protein accumulation by western blot and observed that amino acid substitutions in position 169 did not affect the accumulation of ChpG in mature or premature forms (Figure S7C). This indicate that the observed differences in protein accumulation between ChpG^A^ and ChpG^C^ (Figure S7A, S7B and S7C) is likely to be linked to the signal sequence polymorphic site at position 22 and not to the polymorphic site at position 169. After confirming protein expression, the transformed Cm^101^Ω*chpG* clones were monitored for their ability to elicit HR, cause disease and colonize eggplant. As expected, the V169G substitution in ChpG^A^ abolished its ability to complement Cm^101^Ω*chpG* and bacteria failed to elicit HR and were as pathogenic in eggplant as Cm^101^Ω*chpG* carrying *chpG*^C^ or pHN216 empty vector control (EV) (Figure 7). Correspondingly, the G169V substitution in ChpG^C^ enable it to complement Cm^101^Ω*chpG* and bacteria elicited HR and lost their pathogenicity on eggplant in a similar manner to Cm^101^Ω*chpG* carrying *chpG*^A^ or Cm^101^ (Figure 7). Our analysis confirmed that a single amino acid alteration that occurred in the serine protease domain of ChpG^C^ enable it to evade recognition in eggplant.

**Figure 7.**
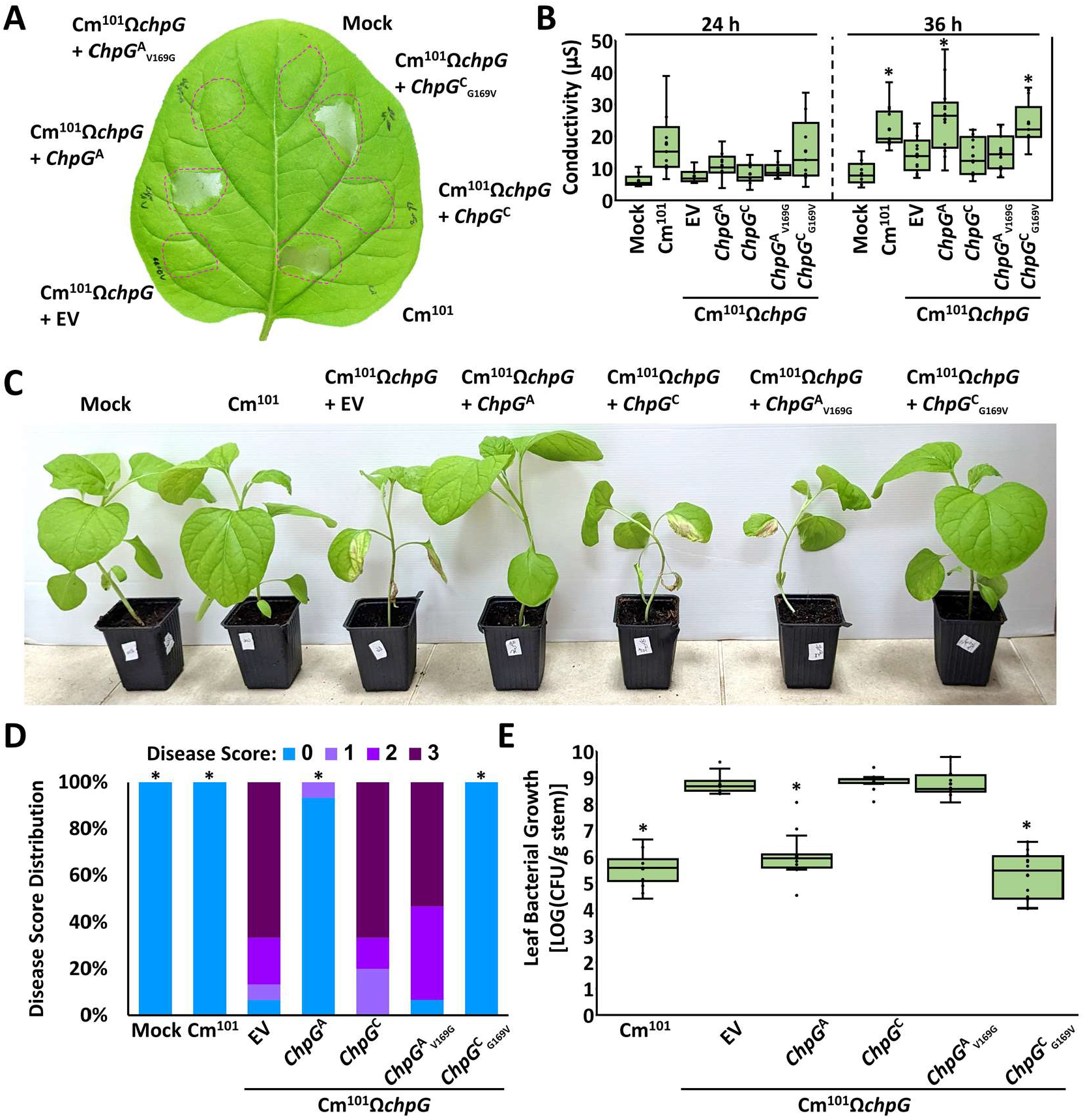
Differential recognition of ChpG variants is linked to a single polymorphic amino acid. (**A, B**) Five to six leaf stage “Black Queen” eggplant leaves were infiltrated (10^8^ CFU/ml) with Cm^101^, Cm^101^Ω*chpG* expressing the indicated *ChpG* variants or empty vector control (EV). (**A**) The picture was taken at 36 hours post inoculation (hpi). (**B**) Cell death was quantified by ion leakage at 24 and 36 hpi. Lower and upper quartiles are marked at the margins of the boxes. Central lines and “o” represent medians and data points of at least 9 (24 h) or 11 (36 h) biological repeats collected from two independent experiments. “*” indicates significant difference (Mann–Whitney U test, *p-value* ≤ 0.05) from Cm^101^Ω*chpG.* (**C, D, E, F**) Three-leaf stage “Black Queen” eggplants were inoculated with the indicated Cm strains or water control (mock) by puncturing the stem area between the cotyledons with a wooden toothpick incubated in *Cm* solution (5 × 10^7^ CFU/ml). (**C, D**) Representative plants were photographed 14 days post inoculations (dpi). (**E**) Leaf blotch symptoms were quantified at 14 dpi according to the following scale: 0 = no leaf blotch, 1 = 1-25%, 2 = 25-50%, 3 = 50-100%. Graph depicts the symptom distribution in at least 15 plants pooled from three independent experiments. “*” indicates the score distribution is different from Cm^101^Ω*chpG* (Pearson’s chi-squared test, *p-value* ≤ 0.05). (**F**) Stem bacterial populations 1 cm above the inoculation sites were quantified at 14 dpi. Lower and upper quartiles are marked at the margins of the boxes. Central lines and “o” represent medians and data points of 16 biological repeats collected from three independent experiments. “*” indicates a significant difference (Mann–Whitney U test, *p-value* ≤ 0.05) from Cm^101^Ω*chpG*.

## Discussion

Plant pathogenic bacteria are subjected to high selective pressures to adapt to their host plants and environment (49). This pressure is potentially increased in seed-borne pathogens that, in addition to constant competition from newly introduced haplotypes, need to adjust to local niches (50). This will dynamically affect local population structures and encourage specialization and establishment of regional adaptive haplotypes that potentially possess different traits than the founder populations (51). Such localized adaptations may challenge the preexisting knowledge of pathogen host range, epidemiology, and pathogenesis mechanisms, which is usually extrapolated from studies that were conducted with representative model strains. In this study, we conducted an in-depth characterization of the Cm population in Israel that combined genomics and virulence-based phenotypic assays. In addition to tomato, we used eggplant in our virulence assays to address the discrepancy in reports regarding whether it is a host of Cm (36)(37)(39)(40). We recently published that eggplant resistance to the Cm model strain Cm^101^ is facilitated through immune recognition of the *chp/tomA* PAI-encoded secreted serine protease ChpG (40). Since most available Cm genomes encode a *chpG* homolog, we hypothesized that eggplant pathogenicity is unique to a Cm subpopulation that was able to evade ChpG recognition. Our findings support this hypothesis and show that while most Cm isolates elicit HR, and are not pathogenic on eggplant; this is not the case for two isolates, C47 and C48. C47 and C48 harbor a unique allelic variant of *chpG* (*chpG*^C^) that is not recognized in eggplant, enabling disease development. This allelic variant differs from the other four *chpG* allelic variants in the Cm population by a single SNP that results in a V169G substitution located in the serine peptidase beta-barrel domain. Intriguingly, we failed to identify the *chpG*^C^ variant in any of the Cm genome deposits available in NCBI, suggesting that this variant is unique to the Israeli Cm population. This is supported by the high phylogenetic proximity between C47 and C48 to other Israeli Cm isolates such as C44 and C46, which do not harbor the V169G substitution (Figure 2A). Therefore, it is more likely that C47 and C48 originated from a parental clone within the Israeli population that acquired this adaptive mutation.

Adaptive mutations and loss in avirulence elicitors have long been hypothesized as the Achilles heel of gene-for-gene-based resistance (52), and host range expansion associated with such adaptations was documented on numerous occasions in plant pathogenic bacteria (53)(54)(55). A recent example can be found in the host range shift of *Xanthomonas euvesicatoria* pv. *perforans*, which was previously considered to be restricted to tomato, but has now extended its host range to pepper due to loss or mutation in the effector *avrBsT* (13)(56). It should be noted that C47 and C48 were originally isolated from tomato. However, we expect host range expansion through minor modification in *chpG* can still harbor potential a beneficial advantage since tomato and eggplant are cultivated in the same geographic regions in Israel and some growers routinely graft eggplants on tomato rootstocks (57)(58). In addition, the host range of these isolates may expend to widespread wild eggplant relatives that potentially recognize ChpG and function as a reservoir for the pathogen such as Buffalobur (*Solanum rostratum*) and Silverleaf (*Solanum elaeagnifolium*) nightshades (59).

A secondary key finding identified by our analyses was the loss of the *chp/tomA* PAI in Cm non-pathogenic isolates. Non-pathogenic *Clavibacter* clones have been previously isolated from tomato in multiple studies. Phylogenetic analyses of these isolates clustered them as either new *Clavibacter* species or as phylogenetically distant Cm strains from a different phyletic lineage of pathogenic Cm strains (60)(61)(62). In contrast to previous findings, phylogenetic analysis of our Cm library clustered all the five identified tomato-non-pathogenic isolates within the Cm lineage. Comparative genomic analysis identified that the *chp/tomA* PAI, which is considered as the major pathogenicity-associated feature of Cm was absent in all five isolates. We initially hypothesized that the five non-pathogenic isolates were progenitor Cm strains that never acquired the *chp/tomA* PAI. However, phylogenetic analysis did not support this hypothesis and showed that all five tomato-non-pathogenic isolates clustered with the pathogenic Cm isolates and can be separated into at least three independent phyletic lineages. This indicated that the *chp/tomA* PAI was more likely to be lost in pathogenic Cm turning them into non-pathogenic strains, and that this event occurred in at least three independent occasions. The *chp/tomA* PAI is flanked by two almost identical direct repeats of ∼1.9 kb (30)(46) which are found in a single copy in the tomato-non-pathogenic isolates, suggesting that that the loss of *chp/tomA* PAI occurred through homologous recombination between these repeats. Looping out of the *chp/tomA* PAI was reported at least twice in the model strain NCPPB382, which resulted in the non-pathogenic strains Cm 30-10 and Cm 27 (30)(15). In both cases, the loss of the *chp/tomA* PAI occurred unintentionally after exposing the bacteria to high voltage during transformation, supporting that such event is feasible and could occur in nature. Considering that the loss of the *chp/tomA* PAI was fixated on multiple occasions suggests that it might be beneficial to Cm in certain conditions. Reduced or abolished virulence due to the loss of large genomic islands associated with pathogenicity has been previously reported in clinical and field isolates of animal and plant pathogenic bacteria such as *Helicobacter pylori*, uropathogenic *Escherichia coli*, *Ralstonia solanacearum* and *Xanthomonas arboricola* (63)(64)(65). Despite its crucial role in growth and pathogenesis in the host, we can speculate on several scenarios that the loss of the *chp/tomA* PAI is beneficial to Cm. The tomato-non-pathogenic isolates in our library originated from tomato plants and can sustain a population of 10^6^-10^7^ CFU/g with very little consequences to the host plant while pathogenic isolates severely damaged the host and, in some occasions, killed the host within weeks after inoculation. Sustaining lower endophytic populations in the host for longer periods might be beneficial to Cm in certain circumstances (66). Alternatively, the presence of the *chp/tomA* PAI might restrict Cm from colonizing alternative hosts through recognition of secreted hydrolases that are specifically recognized in non-host plants such as ChpG or Pat-1 (40)(67)(68). Therefore, losing the *chp/tomA* PAI might expand the number of hosts Cm can occupy as an endophyte. Another possibility is that isolates that lost the *chp/tomA* PAI originated from a cheater sub-population that utilized the *chp/tomA*-encoded secreted hydrolases of the tomato pathogenic isolates as public goods and eventually took over the population (69). Further studies regarding the potential benefits of losing the *chp/tomA* PAI, such as competition assays in host and non-host plants will shed insights into the evolutionary mechanisms that maintain it.

Our study provided novel insights into the phenotypic complexity within the population of bacterial plant pathogens and utilized comparative genomic approaches to link phenotypic variants to distinct genetic features. In addition, we determined that the host range of specialized plant pathogen such as Cm is a variable feature that differs between clones within the population and in our case, is determined by a minor genetic alteration.

## Materials and methods

### Bacterial strains and plant material

*Clavibacter michiganensis* (Cm) isolates used in this study are listed in table S1. *E. coli* strains used in this study are DH5α (Invitrogen) and Rosetta (MERCK) which were utilized for cloning procedures and protein purification, respectively. Cm and *E. coli* were grown in Luria Bertani (LB) broth at 28 or 37°C, respectively. When required, media were supplemented with 10 µg/ml nalidixic acid, 10 µg/ml chloramphenicol, 75 µg/ml neomycin, 50 µg/ml kanamycin or 100 µg/ml ampicillin. Plant cultivars used in this study are tomato (*Solanum lycopersicum*) var. Moneymaker and eggplant (*Solanum melongena*) var. Black Queen. Plants were grown in a 25°C temperature-controlled glasshouse under natural light conditions.

### Assembly of the Cm isolate library

The Cm isolate library consists of 37 isolates which were selected from a collection of more than 250 clones originating from tomato plants displaying bacterial canker symptoms in different regions of Israel from 1994 to 2023 (Table S1). Details regarding the collection area, year of isolation, host type, and any additional information are listed in table S1. Clones that were isolated from 1994 to 2011 were classified into groups according to Pulsed-field gel electrophoresis (PFGE) fingerprinting profile by Dr. Shulamit Manulis-Sasson (21)(42)(Table S1). To further diversify our isolate library, three additional clones (C20, C30, and C31) from a reference Cm strains collection were added as well. These clones were chosen because they had a unique PFGE profile that was different from the other Israeli isolates (Table S1).

### Genome sequencing

Total DNA was extracted from 10 ml bacterial overnight grown cultures using Wizard Genomic DNA Purification Kit (Promega) according to manufacturer’s instructions. Genomic DNA of the Cm isolates was sequenced using an Illumina Nextseq2000 platform with 150bp paired-end reads at the Applied Microbiology Services Laboratory (The Ohio State University). Samples were cleaned using Trimmomatic (70) with default parameters and assembled using Unicycler 0.5.0 and SPAdes v. 3.15.5 (71)(72). Contigs smaller than 200bp were removed, and genome completeness was assessed using BUSCO v5 using micrococcales_odb10 lineage (73).

### Determining *chpG* allelic variants

To determine the presence and allelic variants of *chpG* in our isolates library we conducted a manual Standard Nucleotide BLAST alignment of CMM_0059 against the contigs of all isolates. The aligned DNA sequences of *chpG* ORFs of all isolates and their predicted amino acid were than compared to each other using multiple sequence alignment tool in Clustal Omega (https://www.ebi.ac.uk/Tools/msa/clustalo/). Next, the presence/absence of *chpG* and its allelic variants were confirmed by PCR amplification followed by Sanger sequencing for isolates C3, C4, C5, C6, C18, C21, C29, C30, C31, C45, C46, C47, C48, C61, and C68. To that aim, 1399 bp fragments flanking between the 271 bp upstream to the *chpG* ORF and 294 bp downstream to the *chpG* ORF were amplified with Q5 high fidelity DNA polymerase (NEB) using gene specific primers (Table S3). Each fragment was sequenced by Sanger at Hylabs laboratories using both forward and reverse primer, manually assembled and compared to the *chpG* genomic regions in the corresponding genomes. Specific descriptions of the different *chpG* variants and affiliation with isolates are present in figures S5 and S6 and table 1.

### Plant inoculations, disease severity assessments, and quantification of stem bacterial populations

The virulence assays were carried out using a method similar to that described by Verma and Teper, 2022 (40), with some minor adjustments.

Stem inoculations were conducted by a single punctures of the stem areas between the cotyledons of three-leaf stage eggplants or four-leaf tomatoes with Cm-contaminated toothpicks. Contaminated toothpicks were prepared as follows: Cm bacteria were scraped from fresh 2-day-old cultures grown on LB agar and diluted to 5 × 10^7^ cfu/ml in distilled water in 1.5-ml tubes. Wooden toothpicks were soaked in solutions for at least 10 min and a single toothpick was then used for a single inoculation. After inoculations, plants were kept at 25°C in a glasshouse under natural light conditions and wilt/leaf blotch symptoms were determined and scored at 14 dpi.

Wilting/leaf blotch symptoms were scored in each plant as the percentage of leaves demonstrating wilting and/or necrotic blotch symptoms according to the following scale: 0 = no wilt or leaf blotch, 1 = 1%–25%, 2 = 26%–50%, 3 = 51%–100%.

Bacterial populations were quantified in 1-mm stem areas taken 1 cm above the inoculation sites. Samples were weighed and supplemented with 1 ml of sterile distilled water. Samples were homogenized and bacterial numbers per gram of tissue were determined by plating 10 μl of 10-fold serial dilutions and counting the resulting colonies.

### Plasmid construction and bacterial transformation

Details regarding plasmid construction and primer sequences are available in tables S3 and S4. The pMA-RQ:Cmp plasmid, which was used for sub-cloning of *chpG* variants, was synthesized using GeneArt services (ThermoFisher) and contains a 411 bp fragment composed of the pCMP1 promoter (50302-50044 bp region of CMP1 NCBI GenBank GQ241246) followed by a multiple cloning site and a triple HA tag. To construct *Clavibacter chpG* expression vectors, *chpG* ORFs were amplified from Cm^101^ (Var’ A), CmC5 (Var’ B1), CmC29 (Var’ B2), CmC48 (Var’ C), and CmC6 (Var’ D), and cloned into pMA-RQ:Cmp. For site directed mutagenesis, Val169 of ChpG^A^ and Gly169 of ChpG^C^ were substituted to glycine and valine, respectively, using the QuikChange II kit (Agilent Technologies). The pCMP1:*chpG*:3XHA units were cloned into the *E. coli*-*Clavibacter* shuttle vector pHN216 (45) and transformed into Cm^101^Ω*chpG* and CmC48 as described in Verma and Teper, 2022. Protein accumulation was monitored in lysed overnight Cm cultures by western blot using HA Tag Monoclonal Antibody (2-2.2.14, ThermoFisher) as described by Sambrook and Russell, 2006 (74) and according to the manufacturer’s instructions.

### Expression and purification of MBP fusion proteins in *E. coli*

For construction of *MBP-chpG* fusions plasmids, 112–831 bp fragments containing *chpG* ORFs (CMM_0059) minus the signal peptide-coding region (predicted by SignalP-5.0 Server; http://www.cbs.dtu.dk/services/SignalP/) were amplified from genomic DNA of Cm^101^ and Cm C48, cloned into pMAL-p5x (NEB) and plasmids were introduced into *E. coli* Rosetta cells. Details regarding plasmid construction and primer sequences are available in tables S3 and S4.

For protein purification, bacterial cultures were grown in an orbital shaker at 37°C to OD6_00_ = 0.4–0.6, supplemented with 0.1 mM isopropyl β-d-1-thiogalactopyranoside, and incubated for 4 h at 37°C. Bacteria were pelleted and resuspended in ice-cold buffer solution (20 mM Tris-HCl, 200 mM NaCl, 1 mM EDTA, pH 7.4) and lysed using a SONIC-150W ultrasonic processor (MRC Labs). MBP-fused proteins were purified from supernatants using amylose resin (NEB) according to the manufacturer’s instructions. Purified proteins were quantified by Bradford protein assay kit (Bio-Rad) and validated by SDS-PAGE followed by staining with Coomassie brilliant blue. Protein accumulation in the purified fractions was confirmed with western blot using anti-MBP tag (8G1) mouse monoclonal antibody (Cell Signaling) according to the manufacturer’s instructions.

### Leaf infiltrations and ion leakage measurements

Purified proteins or bacterial suspensions were infiltrated into fully expanded upper rosette leaves of four-to six-leaf stage Black Queen eggplant using a needleless syringe. For protein infiltration, purified MBP, MBP-ChpG^AB^, and MBP-ChpG^C^ proteins were diluted to a concentration of 0.01 µg/ml in 10 mM MgCl_2_ prior to infiltration. For infiltrations with bacterial suspensions, Cm bacteria were scraped from fresh 2-day-old cultures grown on LB agar and diluted to a concentration of 10^8^ (OD_600_ = 0.1) in 10 mM MgCl_2_ prior to infiltration.

For ion leakage measurements, 1.5-cm diameter leaf disks were sampled from the inoculation sites, transferred to flasks containing 10 ml of distilled water, and incubated on an orbital shaker (50 rpm) for 4 h at room temperature. Electrolyte leakage was quantified in water solutions using a conductivity meter (MRC Labs).

## Acknowledgments

We would like to thank Dr. Shulamit Manulis-Sasson (Agricultural Research Organization – Volcani Institute), Ludmila Vagozeb (Israeli Plant Protection and Inspection Services), and Nir Berholtz (Israeli Ministry of Agriculture Extension Service) for providing the Cm isolates used in this study. This work was supported by the Israel Binational Science Foundation (BSF, grant no. 2021190, for D.T. and J.M.J), and the United States-Israel Binational Agricultural Research and Development Fund (BARD; grant no. IS-5499-22, for D.T. and G.L.C).

