## Supplementary material for "Allelic variations in a serine protease effector within *Clavibacter michiganensis* populations determine pathogen host range": Figure S

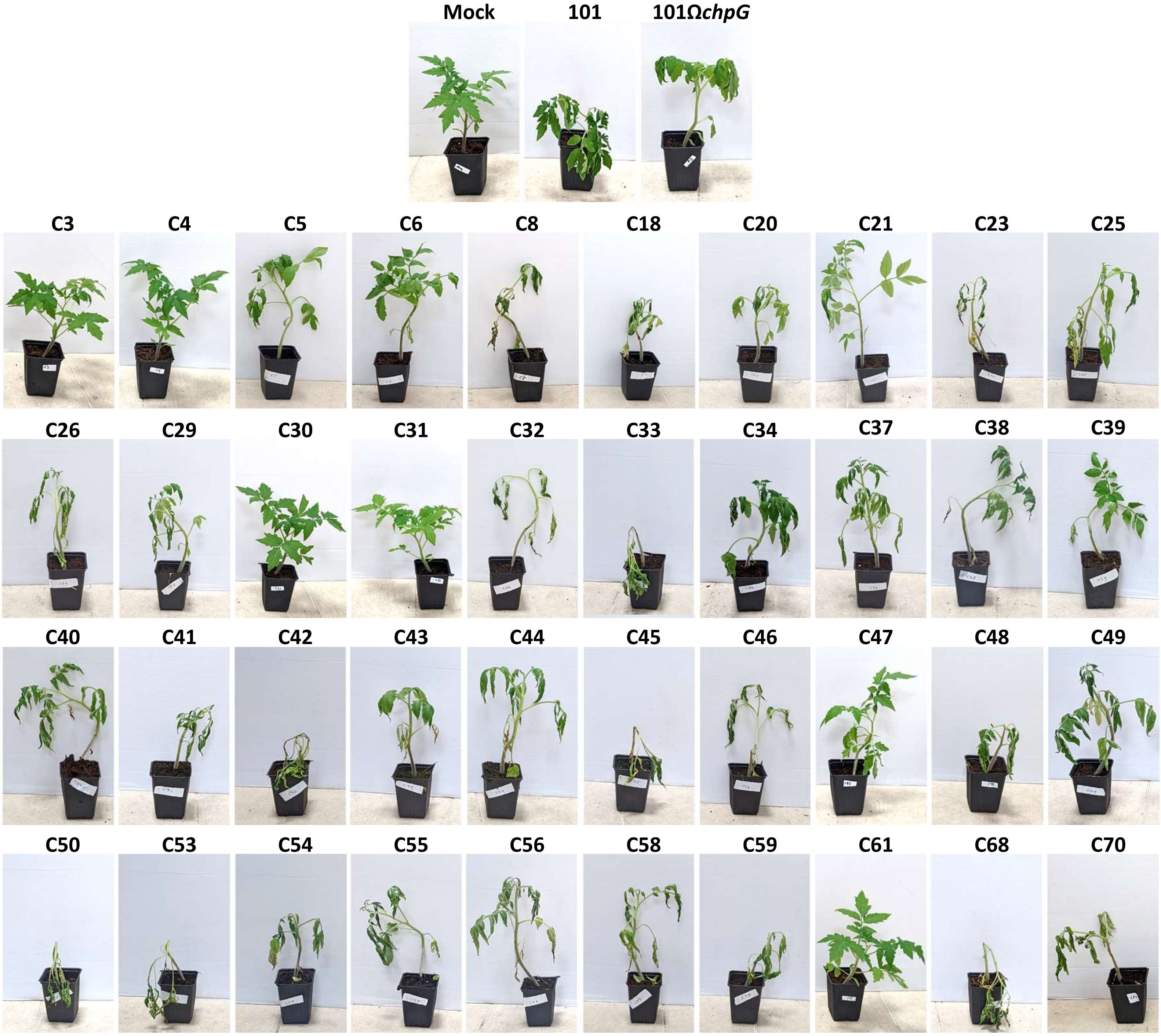

**Figure S1. *Clavibacter michiganensis* (Cm) isolates demonstrate differential virulence in tomato.** Four-leaf stage "Moneymaker" tomato plants were inoculated with the indicated Cm isolates or water control (mock) by puncturing the stem area between the cotyledons with a wooden toothpick incubated in *Cm* solution ($5 \times 10^7$ CFU/ml). Representative plants were photographed 14 days post inoculations. Experiments were repeated at least twice using 3-5 plants for each of the tested Cm isolates.

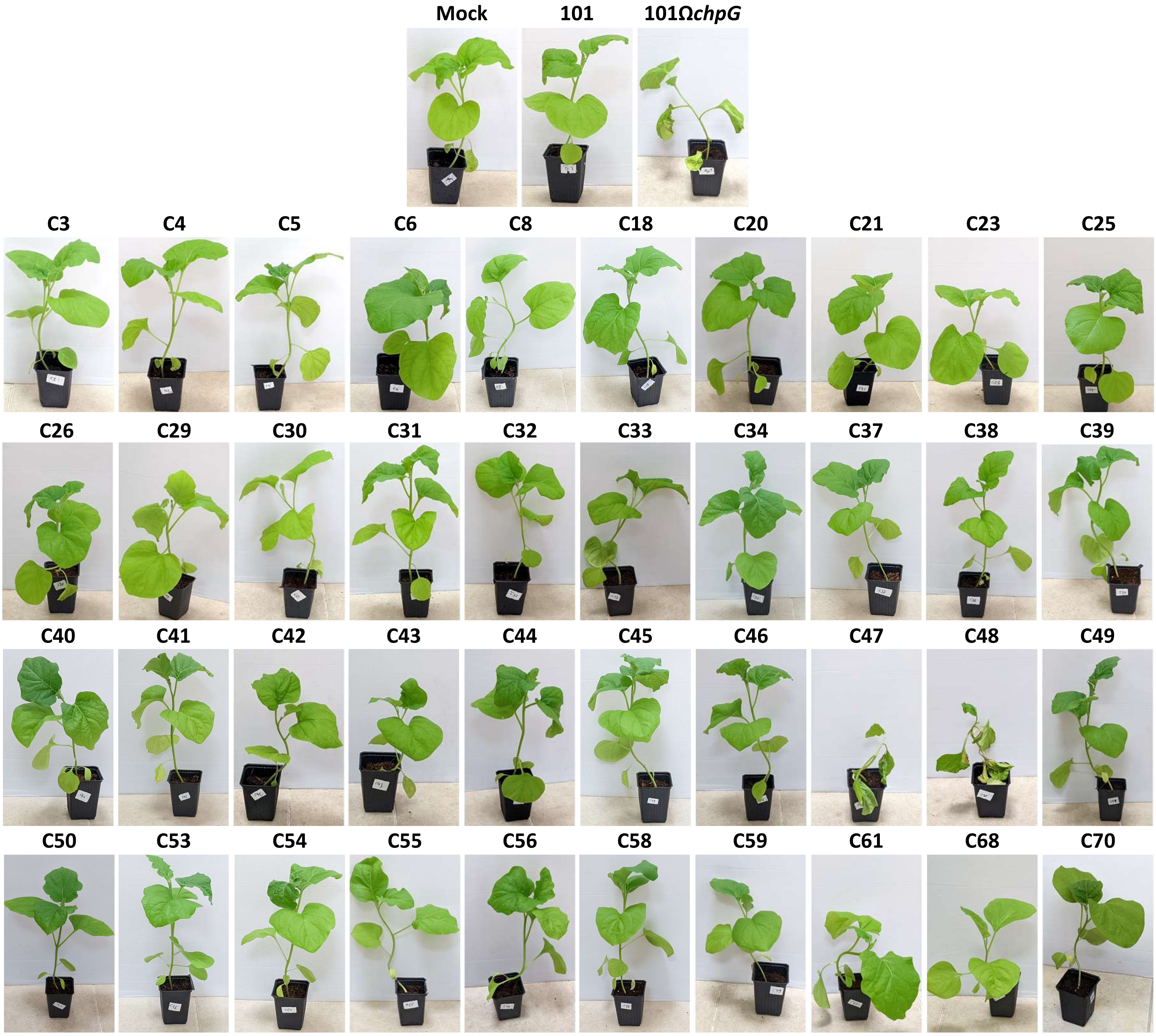

**Figure S2. *Clavibacter michiganensis* (Cm) isolates demonstrate differential virulence in eggplant.** Three-leaf stage "Black Queen" eggplants were inoculated with the indicated *Clavibacter michiganensis* (Cm) isolates or water control (mock) by puncturing the stem area between the cotyledons with a wooden toothpick incubated in *Cm* solution ($5 \times 10^7$ CFU/ml). Representative plants were photographed 14 days post inoculations. Experiments were repeated at least twice using 3-5 plants for each of the tested Cm isolates.

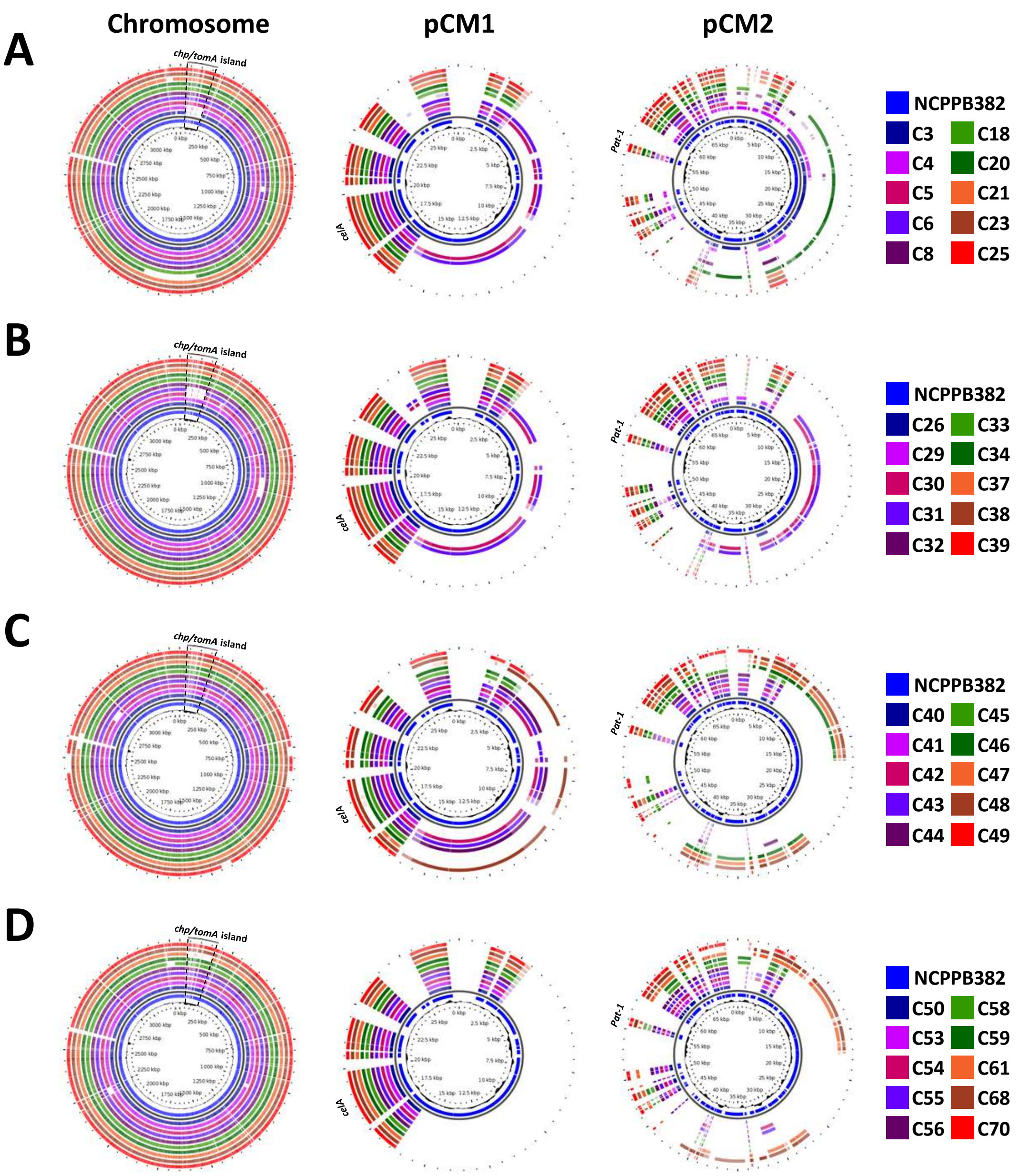

**Figure S3. Genome sequence alignment of tomato-pathogenic *Clavibacter michiganensis* (Cm) isolates**. Whole genome alignment of the tomato pathogenic Cm isolates was done against CDS of Cm strain NCPPB382 chromosome (NCBI GenBank: AM711867), pCM1 plasmid (AM711865), and pCM2 plasmid (AM711866), and was visualized using BLAST atlas analysis in Gview server (https://server.gview.ca/) using default features. The *chp/tomA* island, *celA* (pCM1_0020) and *pat-1* (pCM2_0054) are respectively marked in the chromosome, pCM1, and pCM2 alignments. (**A**) Isolates: C3, C4, C5, C6, C8, C18, C20, C21, C22, C23 and C25. (**B**) Isolates: C26, C29, C30, C31, C32, C33, C34, C37, C38 and C39. (**C**) Isolates: C40, C41, C42, C43, C44, C45, C46, C47, C48 and C49. (**D**) Isolates: C50, C53, C54, C55, C56, C58, C59, C61, C68, and C70.

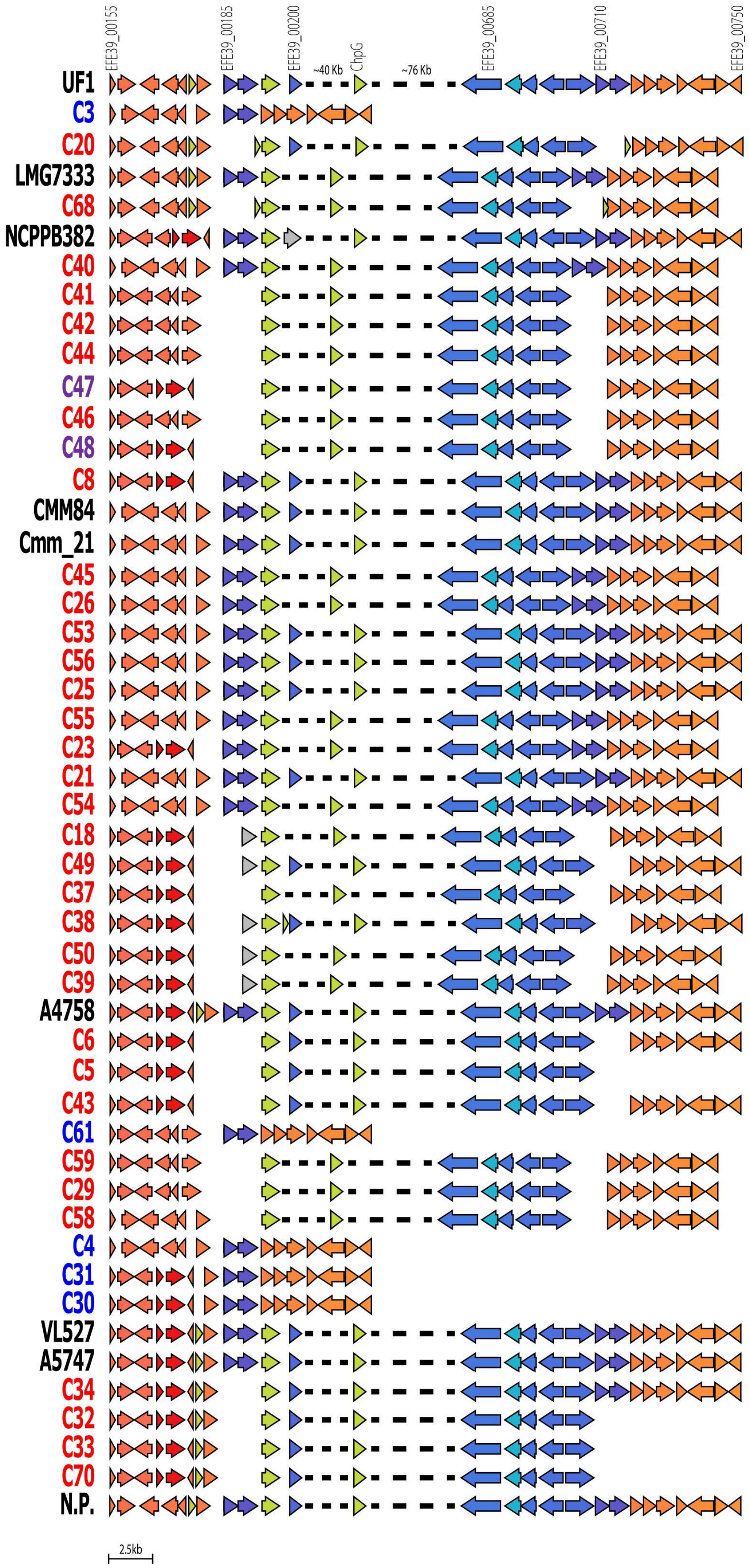

**Fig S4. Global alignment of the genes surrounding the *chp/tomA* island.** The colors indicate homologous gene groups. All coded regions have over 60% alignment sequence identity. Clinker v0.0.27 was used to create the figure, using protein translations predicted by Prokka v1.14.5. Isolates which were pathogenic on tomato but not pathogenic on eggplant are labeled in red, isolates which were pathogenic on tomato and eggplant are labeled in a purple, and isolates which were non-pathogenic on tomato and eggplant are labeled in blue.

```
chpG^A     TTGCCTGCTCGCCATCACACCATCCAGCGCAAGCGCTCAATAGGGGCTGCTCTTCTCGCT   60
chpG^B1    TTGCCTGCTCGCCATCACACCATCCAGCGCAAGCGCTCAATAGGGGCTGCTCTTCTCGCT   60
chpG^B2    TTGCCTGCTCGCCATCACACCATCCAGCGCAAGCGCTCAATAGGGGCTGCTCTTCTCGCT   60
chpG^C     TTGCCTGCTCGCCATCACACCATCCAGCGCAAGCGCTCAATAGGGGCTGCTCTTCTCGCT   60
chpG^D     TTGCCTGCTCGCCATCACACCATCCAGCGCAAGCGCTCAATAGGGGCTGCTCTTCTCGCT   60
           ************************************************************

chpG^A     CTGCCGGCGACCCTTGTCTTGACCTGCATGGCGGGAACACCCGCCTACGCGAACGGACTC   120
chpG^B1    CTGCTGGCGACCCTTGTCTTGACCTGCATGGCGGGAACACCCGCCTACGCGAACGGACTC   120
chpG^B2    CTGCTGGCGACCCTTGTTTTGACCTGCATGGCGGGAACACCCGCCTACGCGAACGGACTC   120
chpG^C     CTGCTGGCGACCCTTGTCTTGACCTGCATGGCGGGAACACCCGCCTACGCGAACGGACTC   120
chpG^D     CTGCTGGCGACCCTTGTCTTGACCTGCATGGCGGGAACACCCGCCTACGCGAACGGACTC   120
           **** ************ ******************************************

chpG^A     AGCAACCCGGACCGCGGAAACTTCCCCATCATCGCCGGTTCCGAAGTCGGCGTTCCGAAT   180
chpG^B1    AGCAACCCGGACCGCGGAAACTTCCCCATCATCGCCGGTTCCGAAGTCGGCGTTCCGAAT   180
chpG^B2    AGCAACCCGGACCGCGGAAACTTCCCCATCATCGCCGGTTCCGAAGTCGGCGTTCCGAAT   180
chpG^C     AGCAACCCGGACCGCGGAAACTTCCCCATCATCGCCGGTTCCGAAGTCGGCGTTCCGAAT   180
chpG^D     AGCAACCCGGACCGCGGAAACTTCCCCATCATCGCCGGTTCCGAAGTCGGCGTTCCGAAT   180
           ************************************************************

chpG^A     GGCTACTGCAGCGTCGGAGCCGTGCTCGTTCCCAGCAGCATCTTCCAGCGGATCACCCCA   240
chpG^B1    GGCTACTGCAGCGTCGGAGCCGTGCTCGTTCCCAGCAGCATCTTCCAGCGGATCACCCCA   240
chpG^B2    GGCTACTGCAGCGTCGGAGCCGTGCTCGTTCCCAGCAGCATCTTCCAGCGGATCACCCCA   240
chpG^C     GGCTACTGCAGCGTCGGAGCCGTGCTCGTTCCCAGCAGCATCTTCCAGCGGATCACCCCA   240
chpG^D     GGCTACTGCAGCGTCGGAGCCGTGCTCGTTCCCAGCAGCATCTTCCAGCGGATCACCCCA   240
           ************************************************************

chpG^A     TATCAGCGCGCTGTTCGCTACCTCGTCCTCGCCAAGCACTGCGCTCCGCTCAACTCGCCC   300
chpG^B1    TATCAGCGCGCTGTTCGCTACCTCGTCCTCGCCAAGCACTGCGCTCCGCTCAACTCGCCC   300
chpG^B2    TATCAGCGCGCTGTTCGCTACCTCGTCCTCGCCAAGCACTGCGCTCCGCTCAACTCGCCC   300
chpG^C     TATCAGCGCGCTGTTCGCTACCTCGTCCTCGCCAAGCACTGCGCTCCGCTCAACTCGCCC   300
chpG^D     TATCAGCGCGCTGTTCGCTACCTCGTCCTCGCCAAGCACTGCGCTCCGCTCAACTCGCCC   300
           ************************************************************

chpG^A     ATCTACTTCGCGCAGCAGGACATCGGAGACGTCGTCTGGCAGTCAGCAGCATCTGACATC   360
```

```
chpG^B1   ATCTACTTCGCGCAGCAGGACATCGGAGACGTCGTCTGGCAGTCAGCAGCATCTGACATC   360
chpG^B2   ATCTACTTCGCGCAGCAGGACATCGGAGACGTCGTCTGGCAGTCAGCAGCATCTGACATC   360
chpG^C    ATCTACTTCGCGCAGCAGGACATCGGAGACGTCGTCTGGCAGTCAGCAGCATCTGACATC   360
chpG^D    ATCTACTTCGCGCAGCAGGACATCGGAGACGTCGTCTGGCAGTCAGCAGCATCTGACATC   360
          ************************************************************

chpG^A    GAGCTGGTCCGCGTATCGCCCTCGCGCGACAACATGACCCTGCACTGCGCCGGCCACTCG   420
chpG^B1   GAGCTGGTCCGCGTATCGCCCTCGCGCGACAACATGACCCTGCACTGCGCCGGCCACTCC   420
chpG^B2   GAGCTGGTCCGCGTATCGCCCTCGCGCGACAACATGACCCTGCACTGCGCCGGCCACTCC   420
chpG^C    GAGCTGGTCCGCGTATCGCCCTCGCGCGACAACATGACCCTGCACTGCGCCGGCCACTCC   420
chpG^D    GAGCTGGTCCGCGTATCGCCCTCGCGCGACAACATGACCCTGCACTGCGCCGGTCACTCC   420
          ***************************************************** *****

chpG^A    AC---CCCCGCAACATGCAGCCCGATCCAGACCTTCACCCCTCGAGCCAACGGCCAAGTC   477
chpG^B1   AC---CCCCGCAACATGCAGCCCGATCCAGACCTTCACCCCTCGAGCCAACGGCCAAGTC   477
chpG^B2   AC---CCCCGCAACATGCAGCCCGATCCAGACCTTCACCCCTCGAGCCAACGGCCAAGTC   477
chpG^C    AC---CCCCGCAACATGCAGCCCGATCCAGACCTTCACCCCTCGAGCCAACGGCCAAGTC   477
chpG^D    ACGAACCCTGCAACATGCAGCCTGATCCAGACCTTCACCCCTCGAGCCAACAGCCAAGTC   480
          **   *** ************** *************************** ********

chpG^A    TTCATGACCGCACCGCCCTCACCGATCGTCGGGCGGCGAGCGATCGCAGGGACGGGTATT   537
chpG^B1   TTCATGACCGCACCGCCCTCACCGATCGTCGGGCGGCGAGCGATCGCAGGGACGGGTATT   537
chpG^B2   TTCATGACCGCACCGCCCTCACCGATCGTCGGGCGGCGAGCGATCGCAGGGACGGGTATT   537
chpG^C    TTCATGACCGCACCGCCCTCACCGATCGGCGGGCGGCGAGCGATCGCAGGGACGGGTATT   537
chpG^D    TTCATGACCGCGCCGCCCTCACCGATCGTCGGGCGGCGAGCGATCGCAGGGACGGGTATT   540
          *********** **************** *******************************

chpG^A    CCGTCGGCCACGGGCACGTTCTGCACGAGCGGGCACGTCACCGGCGTCATCTGCGACTTC   600
chpG^B1   CCGTCGGCCACGGGCACGTTCTGCACGAGCGGGCACGTCACCGGCGTCATCTGCGACTTC   597
chpG^B2   CCGTCGGCCACGGGCACGTTCTGCACGAGCGGGCACGTCACCGGCGTCATCTGCGACTTC   597
chpG^C    CCGTCGGCCACGGGCACGTTCTGCACGAGCGGGCACGTCACCGGCGTCATCTGCGACTTC   597
chpG^D    CCGTCGGCCACGGGCACGTTCTGCACGAGCGGGCACGTCACCGGCGTCATCTGCGACTTC   600
          ************************************************************

chpG^A    CAGCCCACGAGCCTGCCTGTCGGGGTCCTCAGGGCGTATGAGCACCTTGCGGCTGGACAG   657
```

```
chpG^B1     CAGCCCACGAGCCTGCCTGTCGGGGTCCTCAGGGCGTATGAGCACCTTGCGGCTGGACAG    657
chpG^B2     CAGCCCACGAGCCTGCCTGTCGGGGTCCTCAGGGCGTATGAGCACCTTGCGGCTGGACAG    657
chpG^C      CAGCCCACGAGCCTGCCTGTCGGGGTCCTCAGGGCGTATGAGCACCTTGCGGCTGGACAG    657
chpG^D      CAGCCCACGAGCCTGCCTGTCGGGGTCCTCAGGGCGTATGAGCACCTTGCGGCTGGACAG    660
            ************************************************************

chpG^A      TCAGCCGCCGTGGGAGCGCTGCGGCCCGGCGACTCCGGCGGCCCCGTCGTCAGCAAGGAC    717
chpG^B1     TCAGCCGCCGTGGGAGCGCTGCGGCCCGGCGACTCCGGCGGCCCCGTCGTCAGCAAGGAC    717
chpG^B2     TCAGCCGCCGTGGGAGCGCTGCGGCCCGGCGACTCCGGCGGCCCCGTCGTCAGCAAGGAC    717
chpG^C      TCAGCCGCCGTGGGAGCGCTGCGGCCCGGCGACTCCGGCGGCCCCGTCGTCAGCAAGGAC    717
chpG^D      TCAGCCGCCGTGGGAGCGCTGCGGCCCGGCGACTCCGGCGGCCCCGTCGTCAGCAAGGAC    720
            ************************************************************

chpG^A      AGGCGGCTGCTTGGCATCATCTCCGGCGACGTGCCGAACACTCACTTCCTTGTCTACACC    777
chpG^B1     AGGCGGCTGCTTGGCATCATCTCCGGCGACGTGCCGAACACTCACTTCCTTGTCTACACC    777
chpG^B2     AGGCGGCTGCTTGGCATCATCTCCGGCGACGTGCCGAACACTCACTTCCTTGTCTACACC    777
chpG^C      AGGCGGCTGCTTGGCATCATCTCCGGCGACGTGCCGAACACTCACTTCCTTGTCTACACC    777
chpG^D      AGGCGGCTGCTTGGCATCATCTCCGGCGACGTGCCGAACACTCACTTCCTTGTCTACACC    780
            ************************************************************

chpG^A      CCGATGGCGCAGGTCCTCCACGAACTGTCGAGCTACAAGCTCGCACCCGCCAACTGA 834
chpG^B1     CCGATGGCGCAGGTCCTCCACGAACTGTCGAGCTACAAGCTCGCACCCGCCAACTGA 834
chpG^B2     CCGATGGCGCAGGTCCTCCACGAACTGTCGAGCTACAAGCTCGCACCCGCCAACTGA 834
chpG^C      CCGATGGCGCAGGTCCTCCACGAACTGTCGAGCTACAAGCTCGCACCCGCCAACTGA 834
chpG^D      CCGATGGCGCAGGTCCTCCACGAACTGTCGAGCTACAAGCTCGCACCCGCCAACTGA 837
            *********************************************************
```

**Figure S5. DNA sequence alignment of *chpG* homologs.** The five *Clavibacter michiganensis chpG* homologs were aligned by Clustal Omega multiple sequence alignment tool (https://www.ebi.ac.uk/Tools/msa/clustalo/) using default features. Polymorphic site are marked with green (common polymorphic site) or magenta (rare polymorphic site).

```
ChpG^A      MPARHHTIQRKRSIGAALLALPATLVLTCMAGTPAYANGLSNPDRGNFPIIAGSEVGVPN      60
ChpG^B      MPARHHTIQRKRSIGAALLALLATLVLTCMAGTPAYANGLSNPDRGNFPIIAGSEVGVPN      60
ChpG^C      MPARHHTIQRKRSIGAALLALLATLVLTCMAGTPAYANGLSNPDRGNFPIIAGSEVGVPN      60
ChpG^D      MPARHHTIQRKRSIGAALLALLATLVLTCMAGTPAYANGLSNPDRGNFPIIAGSEVGVPN      60
            ********************* **************************************

ChpG^A      GYCSVGAVLVPSSIFQRITPYQRAVRYLVLAKHCAPLNSPIYFAQQDIGDVVWQSAASDI      120
ChpG^B      GYCSVGAVLVPSSIFQRITPYQRAVRYLVLAKHCAPLNSPIYFAQQDIGDVVWQSAASDI      120
ChpG^C      GYCSVGAVLVPSSIFQRITPYQRAVRYLVLAKHCAPLNSPIYFAQQDIGDVVWQSAASDI      120
ChpG^D      GYCSVGAVLVPSSIFQRITPYQRAVRYLVLAKHCAPLNSPIYFAQQDIGDVVWQSAASDI      120
            ************************************************************

ChpG^A      ELVRVSPSRDNMTLHCAGHST-PATCSPIQTFTPRANGQVFMTAPPSPIVGRRAIAGTGI      179
ChpG^B      ELVRVSPSRDNMTLHCAGHST-PATCSPIQTFTPRANGQVFMTAPPSPIVGRRAIAGTGI      179
ChpG^C      ELVRVSPSRDNMTLHCAGHST-PATCSPIQTFTPRANGQVFMTAPPSPIGGRRAIAGTGI      179
ChpG^D      ELVRVSPSRDNMTLHCAGHSTNPATCSLIQTFTPRANSQVFMTAPPSPIVGRRAIAGTGI      180
            ********************* ***** *********.*********** **********

ChpG^A      PSATGTFCTSGHVTGVICDFQPTSLPVGVLRAYEHLAAGQSAAVGALRPGDSGGPVVSKD      239
ChpG^B      PSATGTFCTSGHVTGVICDFQPTSLPVGVLRAYEHLAAGQSAAVGALRPGDSGGPVVSKD      239
ChpG^C      PSATGTFCTSGHVTGVICDFQPTSLPVGVLRAYEHLAAGQSAAVGALRPGDSGGPVVSKD      239
ChpG^D      PSATGTFCTSGHVTGVICDFQPTSLPVGVLRAYEHLAAGQSAAVGALRPGDSGGPVVSKD      240
            ************************************************************

ChpG^A      RRLLGIISGDVPNTHFLVYTPMAQVLHELSSYKLAPAN     277
ChpG^B      RRLLGIISGDVPNTHFLVYTPMAQVLHELSSYKLAPAN     277
ChpG^C      RRLLGIISGDVPNTHFLVYTPMAQVLHELSSYKLAPAN     277
ChpG^D      RRLLGIISGDVPNTHFLVYTPMAQVLHELSSYKLAPAN     278
            **************************************
```

**Figure S6. Amino acid sequence alignment of ChpG homologs.** The four Cm ChpG homologs were aligned by Clustal Omega multiple sequence alignment tool (https://www.ebi.ac.uk/Tools/msa/clustalo/) using default features. Amino acid polymorphic sites are marked with green (common polymorphic site) or magenta (rare polymorphic site).

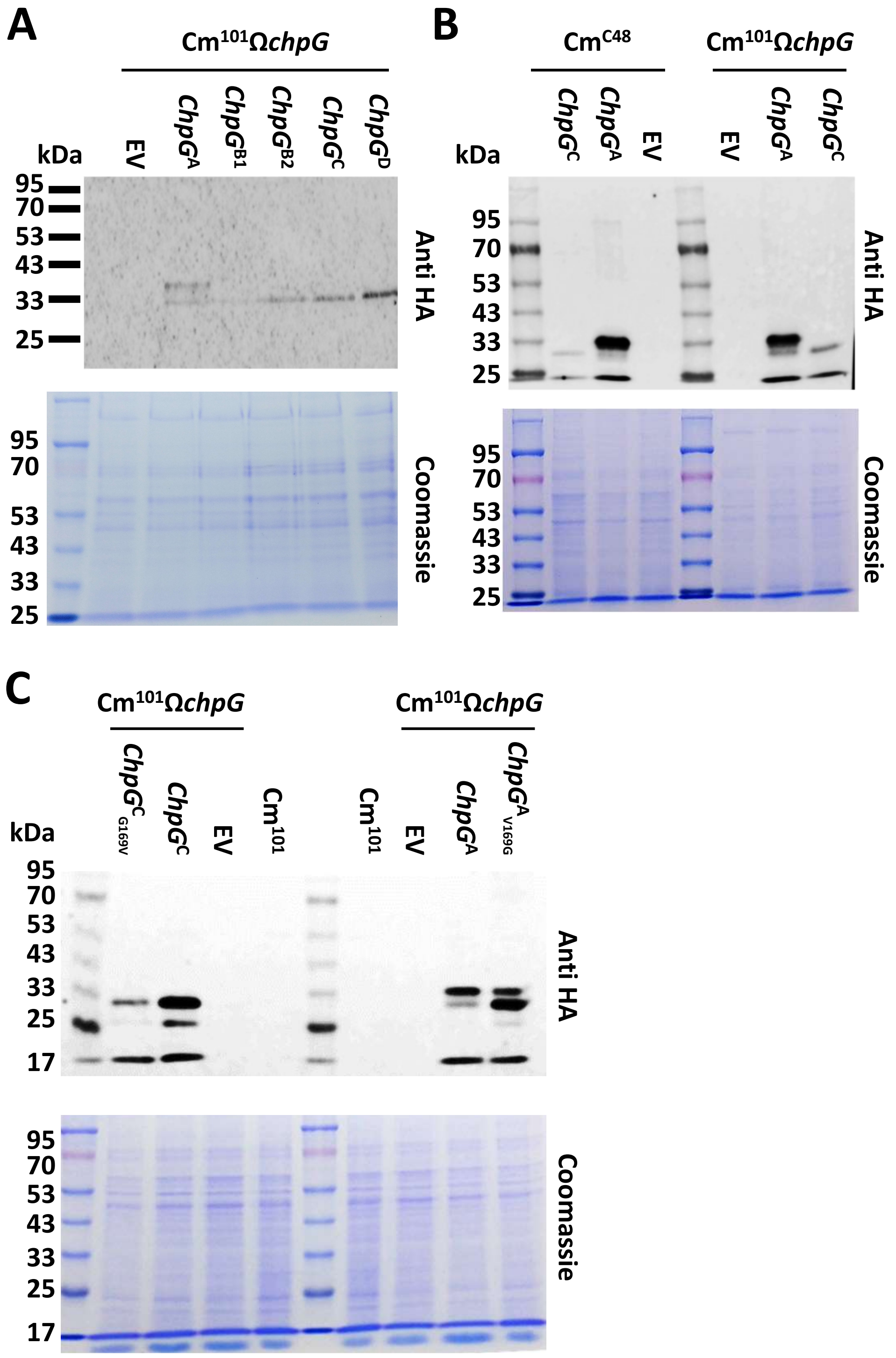

**Figure S7. Protein accumulation of ChpG variants in Cm$^{101}$Ω*chpG* and Cm$^{C48}$.** Overnight cultures of *Clavibacter michiganensis* clones carrying pHN216 with the indicated inserts expressed under the pCMP1 promoter were diluted to OD$_{600}$=0.5, lysed with protein sample buffer and separated on SDS-PAGE and gels were either stained with Coomassie brilliant blue (A, B, C, bottom panel) or transferred to nitrocellulose membrane and immunoblotted with anti-HA antibody (A, B, C, top panel).

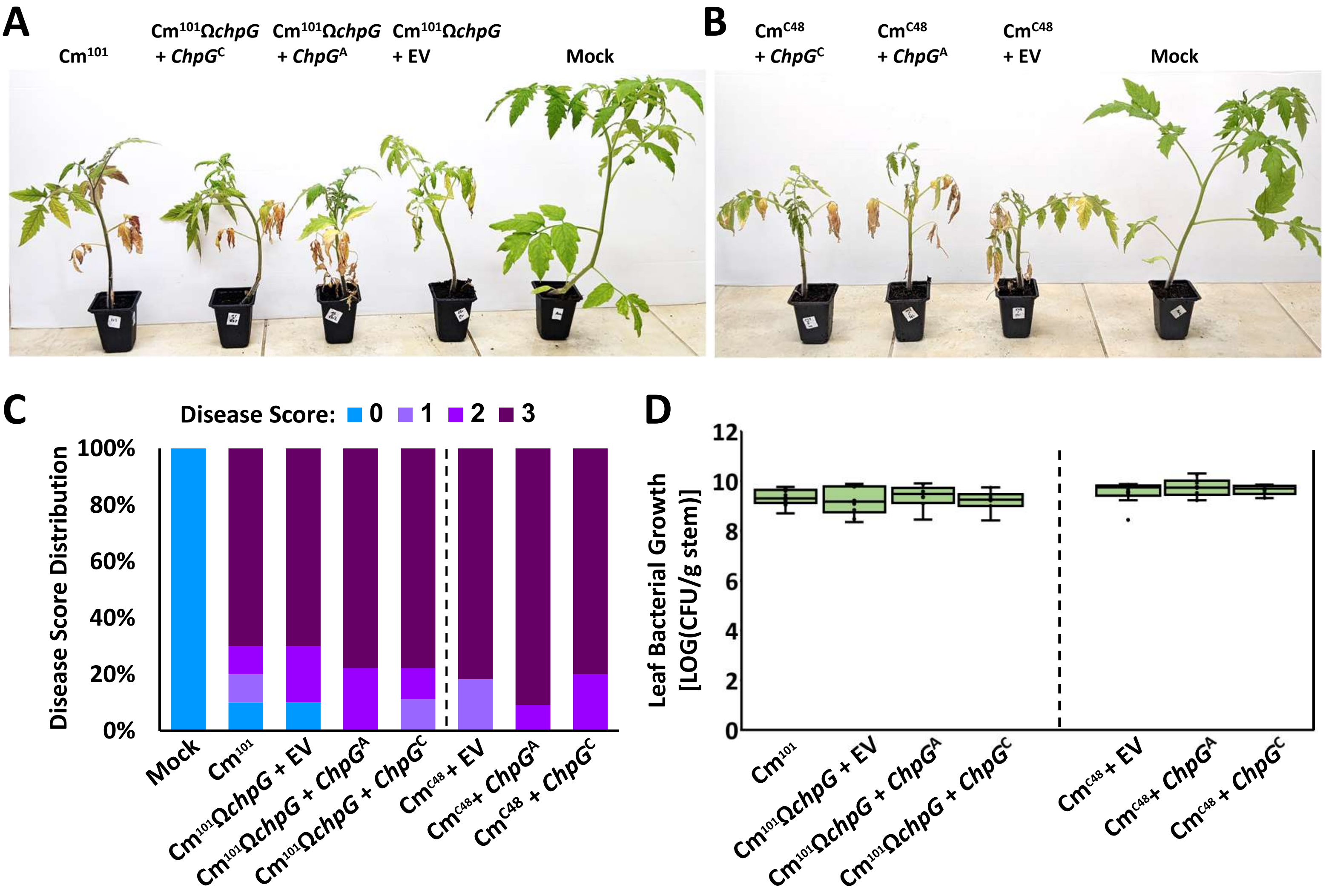

**Figure S8. Introduction of *ChpG*$^{A}$ and *ChpG*$^{C}$ into Cm$^{C48}$ or Cm$^{101}$Ω*chpG* does not affect virulence on tomato**. Four-leaf stage "Moneymaker" tomato plants were inoculated with the indicated *Clavibacter michiganensis* (Cm) clones or water control (mock) by puncturing the stem area between the cotyledons with a wooden toothpick incubated in *Cm* solution ($5 \times 10^7$ CFU/ml). (**A, B**) Representative plants were photographed 21 days post inoculations (dpi). (**C**) Wilting symptoms were quantified at 21 dpi according to the following scale: 0 = no wilting, 1 = 1-25%, 2 = 25-50%, 3 = 50-100%. Graph depicts the symptom distribution in at least nine plants pooled from two independent experiments. (**F**) Stem bacterial populations 1 cm above the inoculation site at 21 dpi. Lower and upper quartiles are marked at the margins of the boxes. Central lines and “o” represent medians and data points of at least nine biological repeats collected from two independent experiments.

**Table S1. Cm clones used in this study**

| Isolate name | Other name | Collection place | Collection region in Israel | Collection year | PFGE cluster | Plant/tissue of isolation | Source | Reference |
|---|---|---|---|---|---|---|---|---|
| Cm101 | CMM101 | Bielefeld University, Germany | NA | 1991 | NT[A] | Lab strain | Rudolf Eichenlaub (Bielefeld University) | (Meletzus et al. 1993) |
| Cm101ΩchpG | CMM101chpGβ | Bielefeld University, Germany | NA | 2008 | NT | Lab strain | Rudolf Eichenlaub (Bielefeld University) | (Stork et al. 2008) |
| C3 | 62 | NA[B] | Gaza strip | 1994 | N | Tomato plant | Shulamit Manulis-Sasson (Volcani Institute) | (Kleitman et al. 2008) |
| C4 | 63 | NA | Jordan valley | 1996 | O | Tomato plant | Shulamit Manulis-Sasson (Volcani Institute) | (Kleitman et al. 2008) |
| C5 | 29 | NA | Gaza strip | 2001 | L | Tomato seeds | Shulamit Manulis-Sasson (Volcani Institute) | (Kleitman et al. 2008) |
| C6 | 30 | Ahitov | Hefer valley | 2001 | M | Tomato fruit | Shulamit Manulis-Sasson (Volcani Institute) | (Kleitman et al. 2008) |
| C8 | 18 | Zipori | Lower Galilee | 1997 | C | Tomato plant | Shulamit Manulis-Sasson (Volcani Institute) | (Kleitman et al. 2008) |
| C18 | 150 | Avni Eithan | Golan heights | 2005 | D | Tomato plant | Shulamit Manulis-Sasson (Volcani Institute) | (Kleitman et al. 2008) |
| C20 | 88 | Netherlands | NA | NA | NC | Tomato plant | Shulamit Manulis-Sasson (Volcani Institute) | (Kleitman et al. 2008) |
| C21 | 32 | Talmi Eliyahu | North-Western Negev | 2001 | B | Tomato plant | Shulamit Manulis-Sasson (Volcani Institute) | (Kleitman et al. 2008) |
| C23 | 64 | NA | Gaza strip | 2000 | B | Tomato plant | Shulamit Manulis-Sasson (Volcani Institute) | (Kleitman et al. 2008) |

| | | | | | | | | |
|---|---|---|---|---|---|---|---|---|
| C25 | 72 | Ami'oz | North-Western Negev | 2002 | B | Tomato plant | Shulamit Manulis-Sasson (Volcani Institute) | (Kleitman et al. 2008) |
| C26 | 110 | Mivtahim | North-Western Negev | 2003 | B | Tomato plant | Shulamit Manulis-Sasson (Volcani Institute) | (Kleitman et al. 2008) |
| C29 | 17 | Prigan | North-Western Negev | 2000 | A | Tomato plant | Shulamit Manulis-Sasson (Volcani Institute) | (Kleitman et al. 2008) |
| C30 | 127 | USA: Michigan | NA | NA | H | Tomato plant | Shulamit Manulis-Sasson (Volcani Institute) | (Kleitman et al. 2008) |
| C31 | 128 | USA: Ohio | NA | NA | H | Tomato plant | Shulamit Manulis-Sasson (Volcani Institute) | (Kleitman et al. 2008) |
| C32 | | Maor | North Coastal Plain | 2019 | NT | Tomato plant | Ludmila Vagozeb (Israeli Plant Protection and Inspection Services) | |
| C33 | | Sde Yitzhak | North Coastal Plain | 2020 | NT | Tomato plant | Ludmila Vagozeb (Israeli Plant Protection and Inspection Services) | |
| C34 | | Sde Yitzhak | North Coastal Plain | 2020 | NT | Tomato plant | Ludmila Vagozeb (Israeli Plant Protection and Inspection Services) | |
| C37 | | Nitzanei Sinai | Central Negev | 2009 | D | Cherry tomato plant | Shulamit Manulis-Sasson (Volcani Institute) | |
| C38 | | Shaked | North Samaria | 2010 | D | Cherry tomato plant | Shulamit Manulis-Sasson (Volcani Institute) | |
| C39 | | Shekef | Lakhish Regional Council | 2010 | D | Tomato plant | Shulamit Manulis-Sasson (Volcani Institute) | |

| | | | | | | | | |
|---|---|---|---|---|---|---|---|---|
| C40 | | Shibolim | North-Western Negev | 2010 | L | Tomato plant | Shulamit Manulis-Sasson (Volcani Institute) | |
| C41 | | Shibolim | North-Western Negev | 2011 | L | Tomato plant | Shulamit Manulis-Sasson (Volcani Institute) | |
| C42 | 551 | Bikaat Arad | Northern Negev | 2011 | L | Tomato plant | Shulamit Manulis-Sasson (Volcani Institute) | (Shtienberg et al. 2018) |
| C43 | 577 | Yesha | North-Western Negev | 2011 | K | Tomato plant | Shulamit Manulis-Sasson (Volcani Institute) | (Shtienberg et al. 2018) |
| C44 | 182 | Hof Hacarmel | North Coastal Plain | 2007 | Z | Tomato plant | Shulamit Manulis-Sasson (Volcani Institute) | (Shtienberg et al. 2018) |
| C45 | 259 | Mivtahim | North-Western Negev | 2009 | Z | Tomato plant | Shulamit Manulis-Sasson (Volcani Institute) | (Shtienberg et al. 2018) |
| C46 | | Ami'oz | North-Western Negev | 2010 | Z | Tomato plant | Shulamit Manulis-Sasson (Volcani Institute) | |
| C47 | | Mlilot | North-Western Negev | 2010 | Z | Tomato plant | Shulamit Manulis-Sasson (Volcani Institute) | |
| C48 | | Netiv HaAsara | North-Western Negev | 2011 | Z | Tomato plant | Shulamit Manulis-Sasson (Volcani Institute) | |
| C49 | 183 | Dekel | North-Western Negev | 2007 | E | NA | Shulamit Manulis-Sasson (Volcani Institute) | (Shtienberg et al. 2018) |
| C50 | | Prigan | North-Western Negev | 2009 | E | Tomato plant | Shulamit Manulis-Sasson (Volcani Institute) | |

| | | | | | | | |
|---|---|---|---|---|---|---|---|
| C53 | | Ein Yahav | Central Negev | 2009 | B | NA | Shulamit Manulis-Sasson (Volcani Institute) |
| C54 | | Ami'oz | North-Western Negev | 2009 | B | Tomato plant | Shulamit Manulis-Sasson (Volcani Institute) |
| C55 | | Yesha | North-Western Negev | 2010 | B | Tomato plant | Shulamit Manulis-Sasson (Volcani Institute) |
| C56 | | Mivtahim | North-Western Negev | 2011 | B | Tomato plant | Shulamit Manulis-Sasson (Volcani Institute) |
| C58 | | Nitzanei Sinai | Central Negev | 2009 | A | Cherry tomato plant | Shulamit Manulis-Sasson (Volcani Institute) |
| C59 | | Mivtahim | North-Western Negev | 2011 | A | Tomato plant | Shulamit Manulis-Sasson (Volcani Institute) |
| C61 | | HaBsor experimental station | North-Western Negev | 2009 | NC | NA | Shulamit Manulis-Sasson (Volcani Institute) |
| C68 | | Shekef | Lakhish Regional Council | 2011 | NC | Cherry tomato plant | Shulamit Manulis-Sasson (Volcani Institute) |
| C70 | | Ami'oz | North-Western Negev | 2023 | NT | Tomato plant | Teper lab (Volcani Institute) |

[A] NT - PFGE analysis was not conducted

[B] NA - Collection year/place is unknown

[C] NC- The isolate does not cluster to any group according to PFGE analysis

**Table S2: Sequencing statistics of Cm isolates sequenced during this study**

| Assembly | Total length | Total contigs | contigs (>= 1000 bp) | Largest contig | GC (%) | N50 | L50 | Accession |
|---|---|---|---|---|---|---|---|---|
| C3 | 3219634 | 27 | 25 | 390595 | 72.87 | 285706 | 5 | SAMN34566069 |
| C4 | 3239786 | 33 | 33 | 385376 | 72.91 | 176568 | 6 | SAMN34566070 |
| C5 | 3295300 | 51 | 43 | 389381 | 72.77 | 152107 | 7 | SAMN34566071 |
| C6 | 3266969 | 54 | 51 | 387924 | 72.8 | 126591 | 8 | SAMN34566072 |
| C8 | 3415064 | 63 | 44 | 555674 | 72.54 | 255269 | 5 | SAMN34566073 |
| C18 | 3346254 | 45 | 41 | 439743 | 72.6 | 258731 | 5 | SAMN34566074 |
| C20 | 3058618 | 50 | 38 | 390541 | 72.52 | 255249 | 5 | SAMN34566076 |
| C21 | 3387424 | 46 | 41 | 328549 | 72.52 | 258771 | 6 | SAMN34566077 |
| C23 | 3422864 | 50 | 44 | 356390 | 72.52 | 150424 | 7 | SAMN34566078 |
| C25 | 3417812 | 44 | 39 | 384951 | 72.52 | 172337 | 6 | SAMN34566079 |
| C26 | 3422263 | 45 | 39 | 458049 | 72.51 | 244537 | 5 | SAMN34566080 |
| C29 | 3296227 | 52 | 48 | 303660 | 72.75 | 110281 | 10 | SAMN34566082 |
| C30 | 3247911 | 28 | 28 | 456785 | 72.76 | 176868 | 6 | SAMN34566083 |
| C31 | 3156240 | 34 | 29 | 461445 | 72.69 | 281461 | 5 | SAMN34566084 |
| C32 | 3296940 | 57 | 48 | 303660 | 72.75 | 110281 | 10 | SAMN34566085 |
| C33 | 3289106 | 66 | 56 | 387672 | 72.79 | 107118 | 10 | SAMN34566086 |

| | | | | | | | | |
|---|---|---|---|---|---|---|---|---|
| C34 | 3318102 | 51 | 37 | 422419 | 72.62 | 301029 | 5 | SAMN34566087 |
| C37 | 3357653 | 56 | 49 | 359398 | 72.56 | 175004 | 7 | SAMN34566090 |
| C38 | 3342280 | 50 | 40 | 388206 | 72.6 | 279224 | 5 | SAMN34566091 |
| C39 | 3347921 | 50 | 45 | 392497 | 72.6 | 134904 | 7 | SAMN34566092 |
| C40 | 3289939 | 48 | 41 | 406300 | 72.79 | 220744 | 5 | SAMN34566093 |
| C41 | 3288693 | 63 | 56 | 387672 | 72.79 | 107118 | 10 | SAMN34566094 |
| C42 | 3288693 | 63 | 56 | 387672 | 72.79 | 107118 | 10 | SAMN34566095 |
| C43 | 3302361 | 42 | 36 | 389694 | 72.75 | 252162 | 5 | SAMN34566096 |
| C44 | 3364495 | 67 | 48 | 422297 | 72.65 | 154368 | 6 | SAMN34566097 |
| C45 | 3347426 | 50 | 41 | 397013 | 72.62 | 297543 | 5 | SAMN34566098 |
| C46 | 3385853 | 51 | 36 | 360537 | 72.59 | 229199 | 6 | SAMN34566099 |
| C47 | 3272850 | 54 | 41 | 360499 | 72.66 | 199457 | 6 | SAMN34566108 |
| C48 | 3385061 | 65 | 44 | 390999 | 72.59 | 259622 | 6 | SAMN34566100 |
| C49 | 3080482 | 88 | 67 | 367749 | 72.49 | 114849 | 8 | SAMN34566101 |
| C50 | 3342280 | 50 | 40 | 388206 | 72.6 | 279224 | 5 | SAMN34566102 |
| C53 | 3317376 | 46 | 37 | 422419 | 72.62 | 301029 | 5 | SAMN34566104 |
| C54 | 3364907 | 42 | 34 | 636165 | 72.56 | 180294 | 5 | SAMN34566105 |
| C55 | 3418633 | 51 | 41 | 576639 | 72.51 | 171727 | 6 | SAMN34566106 |

| | | | | | | | | |
|---|---|---|---|---|---|---|---|---|
| C56 | 3346867 | 54 | 44 | 389053 | 72.62 | 203484 | 6 | SAMN34566107 |
| C58 | 3249338 | 48 | 42 | 354034 | 72.77 | 136390 | 7 | SAMN34566110 |
| C59 | 3292590 | 32 | 32 | 437340 | 72.76 | 218968 | 5 | SAMN34566111 |
| C61 | 3233702 | 43 | 40 | 341864 | 72.92 | 116463 | 8 | SAMN34566112 |
| C68 | 3351655 | 47 | 37 | 400823 | 72.58 | 222040 | 6 | SAMN34566113 |
| C70 | 3398082 | 102 | 46 | 408569 | 72.6 | 153469 | 6 | SAMN34566114 |

**Table S3. Primers used during this study**

| Primer name | Sequence (5’ to 3’) | Destination vector |
|---|---|---|
| *Cloning primers* (bold underline represents restriction sites used for cloning) | | |
| ORFChpGF | AAA**GGATCC**ACGCTACGGAGGAACTCATGC | pMA-RQ:Cmp |
| ORFChpGR | CCC**TCTAGA**GTTGGCGGGTGCCTTGTAG | pMA-RQ:Cmp |
| pMBPChpGF | AA**GGATCC**AACGGACTCAGCAACCCGG | pMALp5x |
| pMBPChpGR | CC**GAATTC**GTTGGCGGGTGCGAGCTTG | pMALp5x |
| *Site directed mutagenesis primers* (bold underline represent nucleotide designated for substitution) | | |
| ChpGT506G-S | ACCGATCG**G**CGGGCGGCGAGCGATCGC | |
| ChpGT506G-AS | GCCGCCCG**C**CGATCGGTGAGGGCGGTG | |
| ChpGG506T-S | ACCGATCG**T**CGGGCGGCGAGCGATCGC | |
| ChpGG506T-AS | GCCGCCCG**A**CGATCGGTGAGGGCGGTG | |
| Amplification/sequencing primers from Cm genomic DNA | | |
| flaChpGF | CTCAGACAACACGACGAGGA | |
| flaChpGR | CTCTCCCTCACCAGACTCCA | |

**Table S4: Plasmids used in this study**

| Plasmid | Relevant characteristic | Reference |
|---|---|---|
| pHN216 | *E. coli-Clavibacter* shuttle vector, based on the replicon of pCM2, $Neo^R/Km^R$ | (Laine *et al.*, 1996) |
| pMA-RQ:Cmp | Synthetic construct carrying a HindII/EcoRI cassette containing pCMP1 promoter followed by a multiple cloning site a triple HA tag. Synthesized by GeneArt service Theromo-Fisher) | This study |
| pHN216:*ChpG*$^A$-3×HA | pHN216 carrying *ChpG*$^A$ (amplified from $Cm^{101}$ strain, locus tag CMM_0059) ORF fused to triple HA-tag under the control of the pCMP1 promoter, $Neo^R/Km^R$ | This study |
| pHN216:*ChpG*$^{B1}$-3×HA | pHN216 carrying *ChpG*$^{B1}$ (amplified from C5 strain, locus tag EFE39_00385) ORF fused to triple HA-tag under the control of the pCMP1 promoter, $Neo^R/Km^R$ | This study |
| pHN216:*ChpG*$^{B2}$-3×HA | pHN216 carrying *ChpG*$^{B2}$ (amplified from C29 strain, locus tag LHJ47_00410) ORF fused to triple HA-tag under the control of the pCMP1 promoter, $Neo^R/Km^R$ | This study |
| pHN216:*ChpG*$^C$-3×HA | pHN216 carrying *ChpG*$^C$ (amplified from Cm C48 strain, locus tag QKF70_15430) ORF fused to triple HA-tag under the control of the pCMP1 promoter, $Neo^R/Km^R$ | This study |
| pHN216:*ChpG*$^D$-3×HA | pHN216 carrying *ChpG*$^D$ (amplified from C6 strain, locus tag QKG63_14905) ORF fused to triple HA-tag under the control of the pCMP1 promoter, $Neo^R/Km^R$ | This study |
| pHN216:*ChpG*$^A_{V169G}$-3×HA | pHN216:*ChpG* carrying a T506->G substitution within the ORF of *ChpG*$^A$ (CMM_0059), $Neo^R/Km^R$ | This study |
| pHN216:*ChpG*$^C_{G169V}$-3×HA | pHN216:*ChpG* carrying a G506->T substitution within the ORF of *ChpG*$^C$, $Neo^R/Km^R$ | This study |
| pMALp5x | *E. coli* expression vector containing secreted maltose binding protein (MBP) tag, $Amp^R$ | NEB (Ipswich,MA, USA) |
| pMALp5x:*ChpG*$^{A/B}$ | pMALp5x carrying the 112-831 bp fragment (ORF minus signal peptide coding region) of *ChpG*$^A$ or *ChpG*$^B$ introduced into the BamHI/EcoRI sites, $Amp^R$ | This study |
| pMALp5x:*ChpG*$^C$ | pMALp5x carrying the 112-831 bp fragment (ORF minus signal peptide coding region) of *ChpG*$^C$ introduced into the BamHI/EcoRI sites, $Amp^R$ | This study |

*$Neo^R$, $Km^R$ and $Amp^R$ indicate chloramphenicol, neomycin, kanamycin and ampicillin resistance, respectively
